# Gene regulatory co-option drives birdsong neural circuit specialization

**DOI:** 10.64898/2026.04.24.720702

**Authors:** Arturo P. Marquez, Cayla R. M. Lagousis, Eva G. Katcher, Frank Rand, Keerthi Varadharaj, Michael S. Brainard, Bradley M. Colquitt

## Abstract

As animals evolve complex motor skills, they acquire more diverse supporting motor circuits in their nervous systems. Yet the molecular mechanisms driving motor circuit evolution remain poorly understood. Birdsong, a learned complex motor skill with parallels to human speech, is controlled by a dedicated neural circuit – the song system – that is distinguished from nearby sensorimotor regions by molecular, physiological, and connectivity specializations. By profiling gene expression and chromatin accessibility in the songbird brain, we found that each projection neuron type in the song system has a molecularly similar sister neuron type in adjacent non-song regions; these sister neurons lack specialized gene expression and are transcriptionally similar to neurons in the chicken brain. The gene regulatory networks (GRNs) controlled by transcription factors *MAFB* and *EMX2*, typically active in fast-spiking interneurons and astrocytes, are specifically active in song-dedicated extratelencephalic projection neurons. Furthermore, the heterologous expression of *MAFB* or *EMX2* in chicken projection neurons was sufficient to drive expression programs characteristic of song neurons. These results support a model in which song-dedicated neurons emerged from ancestral neural types in part through the co-option of GRNs active in other cellular contexts, providing a genetic mechanism underlying the evolution of birdsong.

## Introduction

The complexity of motor skills, such as forelimb dexterity and human speech, is correlated with an expansion and diversification of motor cortical domains [1]. Primates with fine control of the hands have a primary motor cortex (M1) consisting of an evolutionarily “old” domain present in all primates and a “new” domain defined by the presence of corticospinal neurons that make direct connections to motoneurons in the ventral horn [2]. Similarly, the human brain contains two motor regions that control the larynx, a ventral domain present across primates and a dorsal domain found only in humans [3]. The ability to perform advanced motor skills relies on these specialized neural regions and cell types due to their specific molecular, physiological, and connectivity properties [1, 2, 4]. These examples support a general framework in which novel behavioral capacities evolve through the repurposing and refining of ancestral neural architectures. However, the molecular mechanisms that drive innovations in motor circuits remain poorly characterized.

Imitative vocal learning is a complex motor skill that has evolved repeatedly in several independent mammalian and avian lineages [5]. Among these lineages, songbirds offer an experimentally tractable system for investigating the molecular and circuit-level mechanisms that support the evolution of motor skills. Birdsong is generated by a dedicated neural circuit – the song system – consisting of multiple interconnected brain regions that are embedded within, but anatomically distinct from, sensorimotor areas that control other behaviors. Birds that do not learn their vocalizations largely lack these specialized regions, supporting a model in which the song system is a derived neural circuit that evolved from ancestral avian sensorimotor circuits (**Fig. 1a**) [6, 7]. These features make birdsong and its associated circuitry a powerful model for identifying the molecular determinants of motor circuit specialization and the mechanisms influencing the evolution of motor skills.

**Figure 1:**
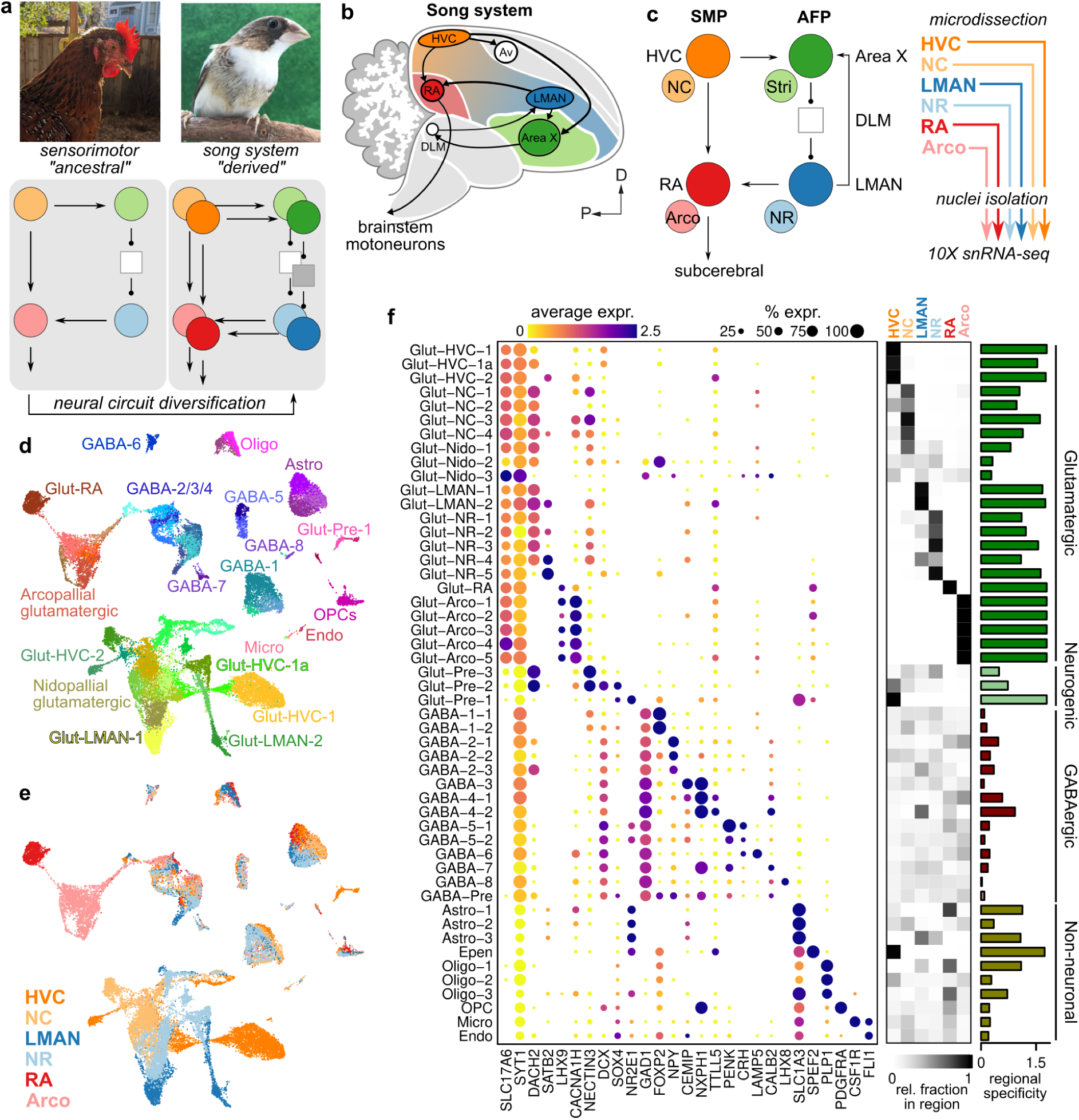
Single-nucleus RNA-sequencing of birdsong-dedicated brain regions and surrounding non-song regions. **(a)** Model of the evolution of the song system, the neural circuit controlling birdsong, from ancestral sensorimotor circuits. **(b)** Schematic overview of the song system. HVC, proper name; RA, robust nucleus of the arcopallium; LMAN, lateral magnocellular nucleus of the anterior nidopallium; Av, avalanche; DLM, medial portion of the dorsolateral thalamic nucleus; D, dorsal; P, posterior. **(c)** Circuit diagram of the song system and sample collection. Arrowheads and closed circles indicate excitatory and inhibitory connections, respectively. NC, caudal nidopallium; Arco., arcopallium; NR, rostral nidopallium; Stri., striatum. **(d)** Uniform manifold approximate projection (UMAP) plots of snRNA-seq data generated from HVC, LMAN, RA, and surrounding non-song regions. Points colored by cell cluster. **(e)** As panel **d** with points colored by region. **(f)** *(Left*) Expression of marker genes across clusters. (*Right*) Comparison of the regional distributions of each cell type. Heatmap intensity indicates the relative proportion of a given cell type in each region. Bar graph at right indicates the region specificity (see Methods) of each cell type.

Here, we tested the hypothesis that song-dedicated neurons are a product of cellular evolution and that these neurons engage gene regulatory networks that drive their specialization away from neurons in adjacent regions. To do so, we characterized the transcriptional and gene regulatory differences between song and non-song motor regions of a model songbird brain (the Bengalese finch, *Lonchura striata domestica*) using single-nucleus RNA and ATAC sequencing, gene regulatory network inference, comparative genomics, and cross-species gene manipulations. We found that song glutamatergic neurons exhibit transcriptional signatures consistent with their evolution from ancestral sensorimotor neuron classes present across avian brains, with each song neuron type having a paired non-song sister cell type from which it diverged. This divergence is driven in part via the co-option of the gene regulatory networks for *MAFB* and *EMX2*. Overexpressing either of these transcription factors in chicken telencephalons is sufficient to partially induce a song neuron-like expression profile in chicken glutamatergic neurons, indicating that these factors can act as drivers of song circuit specialization even across substantial evolutionary distance.

## Results

### Transcriptional specializations of the song system

The song system consists of two functional pathways, the motor pathway and the anterior forebrain pathway (AFP) that work in a coordinated manner to enable the learning and production of birdsong (**Fig. 1b,c**). The motor pathway, made up of RA (robust nucleus of the arcopallium) and HVC (used as a proper noun), is responsible for the patterning and coordination of breathing and vocal muscle activity in song production [8–11]. In contrast, the AFP, consisting of LMAN (lateral magnocellular nucleus of the anterior nidopallium), Area X, and DLM (medial nucleus of the dorsolateral thalamus), is a specialized basal ganglia thalamocortical loop that plays a significant role during song learning in juveniles and adaptive plasticity in adults [12–16].

Song regions display a number of transcriptional, physiological, and connectivity differences compared to surrounding areas that likely underpin the distinct features of birdsong relative to other motor behaviors [17–19]. However, a comprehensive comparison of the molecular differences between the individual cell types in the song system versus those in surrounding non-song areas has not been performed. To characterize these transcriptional differences, we performed single-nucleus RNA-seq on microdissected song regions – RA, HVC, LMAN – and adjacent non-song regions – arcopallium, caudal nidopallium, and rostral nidopallium (**Fig. 1c** and **Extended Data Fig. 1a,b**).

As in our previous study [20], we found that glutamatergic neurons (”Glut”) differ substantially at the transcriptional level between regions, such that each type of Glut neuron is found in only one to four regions (**Fig. 1d-f**). Cell type clusters were defined on the full set of glutamatergic neurons then assigned labels according to their distributions across the six microdissected regions. HVC samples contained two distinct cell types that were found exclusively in HVC (Glut-HVC-1 and Glut-HVC-2) (**Fig. 1f**), with a third (Glut-HVC-1a) that was intermediate between Glut-HVC-1 and nidopallial regions outside of HVC (**Fig. 1d**). Similarly, LMAN samples contained two exclusive projection neuron types (Glut-LMAN-1, Glut-LMAN-2) and RA a single type (Glut-RA). Outside of song regions, we found several projection neuron types located predominantly in the caudal nidopallium (”NC”), rostral nidopallium (”NR”), spanning multiple nidopallial regions (”Nido”), and the arcopallium (”Arco”) (see Supplementary Discussion for further interpretation of cell type distributions).

In contrast, each type of GABAergic neuron (”GABA”) and non-neuronal cell is distributed in largely equal proportions across brain regions. We found eight classes of GABAergic neurons (GABA-1 through GABA-8), which matched the classes we previously identified in our analysis of only HVC and RA (**Fig. 1d,f**). These classes were similarly distributed between each song region and its adjacent non-song region. However, there were some exceptions to this even distribution, including a distinct population of *Pvalb*-class interneurons (GABA-4-2) that is enriched in LMAN and RA and subclasses of *Npy*-class interneurons (GABA-2) that are differentially distributed across regions (nidopallium versus arcopallium) (**Fig. 1f**).

We identified three populations of astrocytes: one primarily found in the song regions HVC and RA (Astro-1), a caudal population present in HVC, NC, RA, and Arco (Astro-2), and a rostral population enriched in LMAN and NR (Astro-3) (**Fig. 1f**). We also detected three populations of oligodendrocytes that were each enriched in song regions relative to non-song regions (**Fig. 1f**), consistent with the high levels of myelination in the song system [21].

### Song glutamatergic neurons have paired counterparts in surrounding regions

We hypothesized that the transcriptional differences between glutamatergic neurons in song regions and adjacent regions reflect the specialization of song-dedicated glutamatergic neurons from a more generalist glutamatergic type present in non-song regions. We reasoned that we could gain insight into the “baseline” expression profile of song glutamatergic neurons by computationally removing genes that are differentially enriched in these neurons relative to other glutamatergic neurons (see Methods).

Consistent with the specialization model, we found that after removing genes with song-specialized expression, each song glutamatergic neuron clustered with a non-song type (**Fig. 2a,b**). Glut-RA clustered with Glut-Arco-1, which displays expression patterns similar to the arcopallial motor region (dorsal intermediate arcopallium) (**Extended Data Fig. 2a,b**). Glut-HVC-1 was paired with Glut-NC-1, while Glut-HVC-2 was paired with Glut-NC-4. LMAN, the other song region in the nidopallium, displayed a similar pattern: Glut-LMAN-1 clustered with Glut-NR-1 and Glut-LMAN-2 clustered with Glut-NR-4. Furthermore, the two principal projection neuron classes in HVC and LMAN clustered with each other (Glut-HVC-1 with Glut-LMAN-1 and Glut-HVC-2 with Glut-LMAN-2), suggesting that each region contains a similar set of projection neurons. In our previous work [20], we used retrograde tracing to determine that Glut-HVC-1 neurons project to RA (HVC_RA_, “corticocortical”) while Glut-HVC-2 neurons project to Area X (HVC_X_, “corticostriatal”). Like HVC, LMAN also projects to both RA and Area X, suggesting that its two projection classes have similar connectivities (however, see **Extended Data Fig. 2c** and Supplementary Discussion for discussion of cluster labeling).

**Figure 2:**
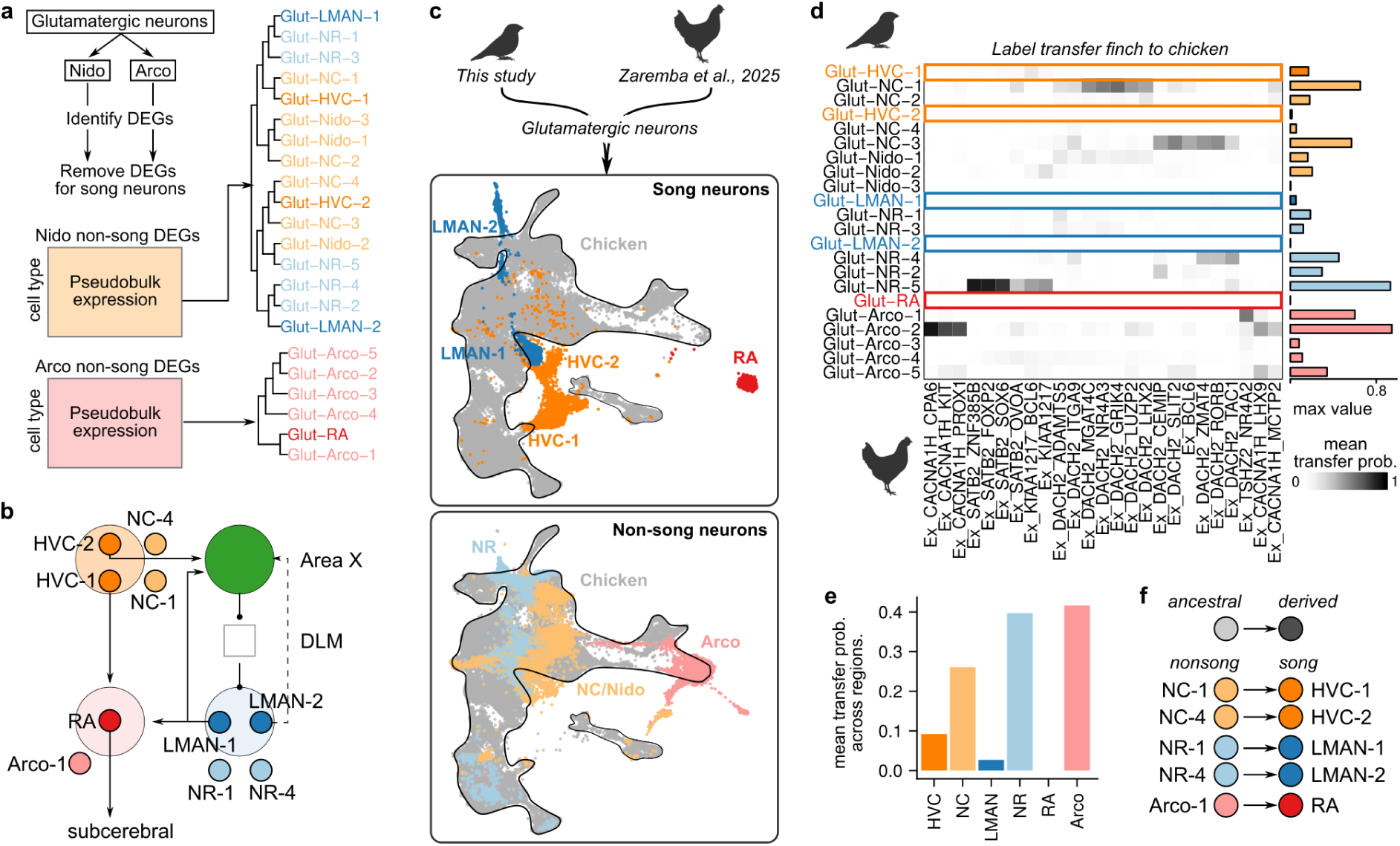
Transcriptional divergence of song glutamatergic neurons from non-song neurons in the songbird and chicken brain. **(a)** Hierarchical clustering of differentially expressed genes across glutamatergic neurons in the nidopallium and arcopallium. Genes that were differentially expressed in song neurons were removed before clustering to assess non-specialized patterns of transcriptional similarity. **(b)** Circuit model of pallial song projection neurons paired with neuron types in non-song regions. **(c)** Integration of snRNA-seq data from songbird (this study) and chicken (Zaremba et al. 2025) glutamatergic neurons via reciprocal PCA (RCPA). UMAPs show a single joint embedding of songbird and chicken cells, with either (*top*) song neurons or (*bottom*) non-song neurons displayed with chicken glutamatergic neurons (gray, out-lined) **(d)** Heatmap intensity indicates the mean label transfer probability between each neuron pair. The bar plot at right of the heatmap represents the maximum label transfer probability for each songbird neuron type. **(e)** Average transfer probability across songbird glutamatergic neurons separated by brain region. **(f)** Model of specialization. Distinct sets of glutamatergic neurons outside of the song system represent ancestral cell states that are similar to those found in the corresponding brain regions of non-vocal learning species. Specialized projection neurons in song regions represent a derived cell state found specifically in the songbird clade.

To determine whether glutamatergic neurons in the separate regions of the song system exhibit common or distinct specialization programs from complementary non-song neurons, we analyzed differential expression between each pair of song and non-song glutamatergic neuron type (**Extended Data Fig. 3a-c**). Although each song projection neuron type exhibited a distinct transcriptional signature of specialization, nidopallial song neurons showed greater similarities in their specialization programs with each other relative to arcopallial Glut-RA vs. Glut-Arco-1 (**Extended Data Fig. 3c,d**). The specialization programs for each song neuron were enriched for similar ontology terms, including a reduction in synapse and synaptic plasticity related genes and an increase in extracellular matrix associated genes (**Extended Data Fig. 3e**). Together, these expression profiles reflect the synaptic and structural stability of the song system in adult male finches, in keeping with the high performance consistency of adult birdsong.

### Song glutamatergic neurons specialize from ancestral non-song neurons

One model of song system evolution proposes that the circuit diverged from an ancestral sensorimotor circuit to acquire the specialized properties required to support the motor and imitative learning demands of birdsong [6, 22]. A prediction of this model is that neurons in the non-specialized sensorimotor regions of songbirds have similar properties to those found in sensorimotor regions of birds that do not have complex, learned vocal repertoires, such as chickens. This prediction suggests that non-specialized sensorimotor neurons in songbirds and sensorimotor neurons in non-vocal learning birds represent ancestral neuronal states, while song-specialized neurons represent a derived neuronal cell type in the songbird clade (**Fig. 1a**). To test this prediction, we integrated our snRNA-seq dataset with a recently published snRNA-seq dataset of the adult chicken brain [23] and analyzed (1) whether song-specialized glutamatergic neurons represent distinct neuron types that are not found in the chicken brain and (2) whether non-song-specialized glutamatergic neurons overlap with populations of neurons in the chicken brain (**Fig. 2c-f**).

We found that neuron types in non-song regions demonstrated transcriptional similarity with glutamatergic neuron types in chickens (**Fig. 2c-e**). In contrast, each song neuron class did not integrate well with any glutamatergic neuron class in the chicken telencephalon. Together, these results support the model in which song projection neurons specialize from sensorimotor neuron classes that are present across avian brains (**Fig. 2f**). We propose that these relationships are the end result of cellular evolution within the songbird clade: non-song projection neurons represent “ancestral” classes, while song-specialized projection neurons represent “derived” classes present only in the songbird lineage. Thus, each song neuron and its paired non-song neuron class can be considered sister cell types [24], and identifying the regulatory mechanisms that drive their divergence will provide insight into the molecular control of birdsong neural circuit specialization.

### Identification of song-specialized gene regulatory networks

Gene regulatory networks (GRNs) generate diverse cell types during development and their evolution can generate cellular innovation. The extensive transcriptional differences between each song projection neuron and its non-song neuron type suggest that distinct gene regulatory programs drive song neuron divergence. To infer these networks, we performed single-nucleus RNA and ATAC-seq (10X Genomics Multiome-seq) on microdissected HVC, RA, NC, and Arco (**Fig. 3a,b** and **Extended Data Fig. 4a**).

**Figure 3:**
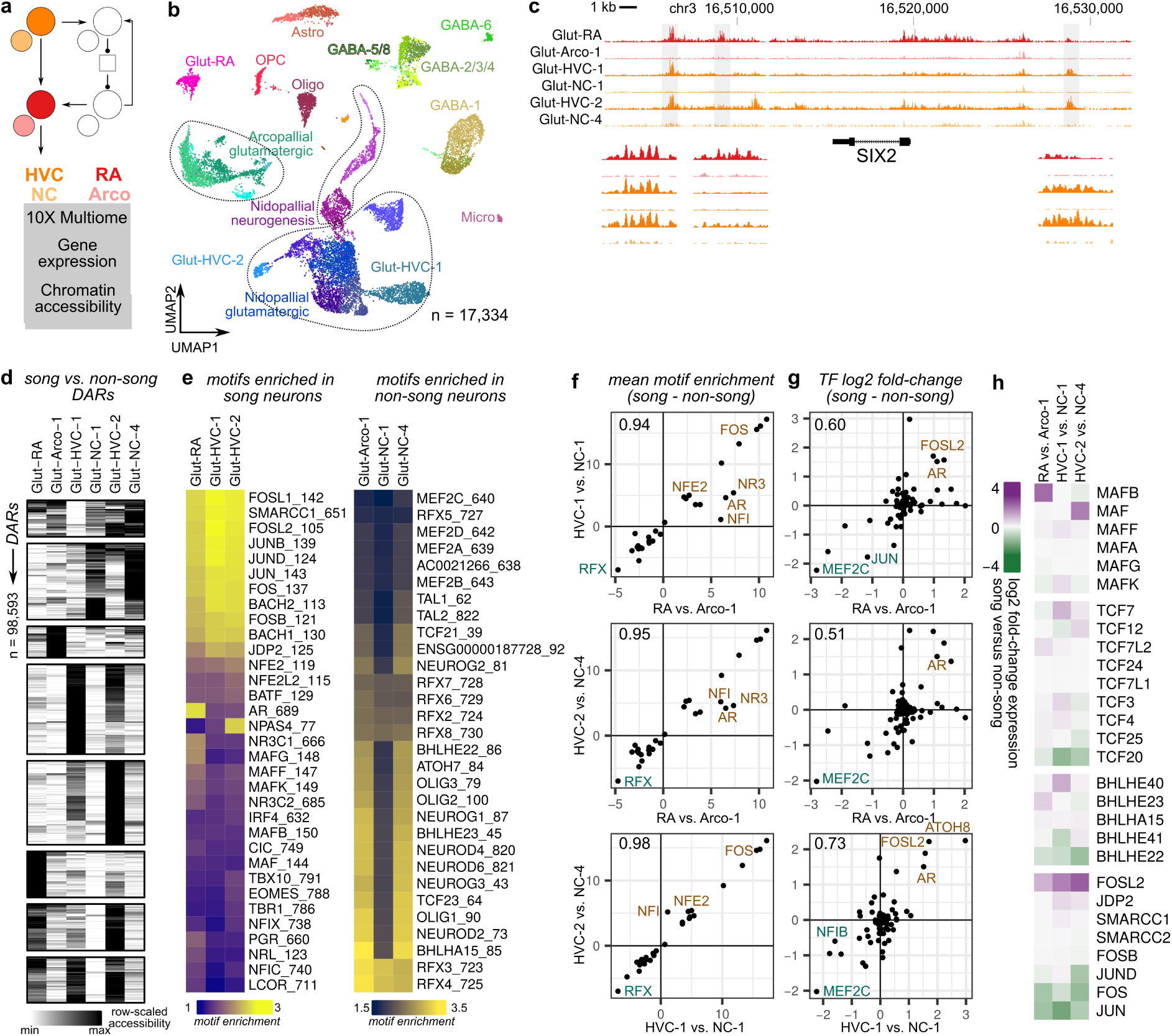
Shared and cell type-specific gene regulatory programs underlie song neu-ron divergence. **(a)** 10X Multiome-seq was performed on two regions of the song motor pathway, HVC and RA, along with adjacent non-song regions, NC and Arco. **(b)** UMAP plot of the Multiome dataset indicating cluster labels **(c)** Chromatin accessibility at the *SIX2* locus, a transcription factor with elevated expression in Glut-RA and Glut-HVC-2 relative to their paired non-song neurons Glut-Arco-1 and Glut-NC-4. **(d)** Accessibility across song and paired non-song glutamatergic neurons in differentially accessible regions (DARs), separated according to k-means clustering. Shown is a random subset (n=1,000) of the total number of DARs. **(e)** Transcription factor binding motifs enriched in DARs with higher accessiblity in (*left*) song vs. non-song neurons or (*right*) non-song vs. song neurons. **(f)** Shared patterns of differential motif enrichment in DARs between song and non-song neurons. The value in the top left of each box is Pearson’s *r*. **(g)** Transcription factor differential expression between song and non-song neurons exhibits less consistency than motifs across song glutamatergic neurons. **(h)** Differential expression of several families of transcription factors (MAF, TCF, BHLH, and AP-1 complex).

To determine how accessibility profiles varied between song and non-song regions across cell types, we identified differentially accessible regions (DARs) within broad cell classes located in either HVC or Nido and RA or Arco. Similar to gene expression variation across cell types, glutamatergic neurons exhibited the highest differences in accessibility profiles between song and non-song regions, while those of GABAergic neurons and non-neuronal cells showed little difference between regions (**Extended Data Fig. 4b**). We then identified DARs between each song glutamatergic neuron type and its partner in the adjacent non-song area: Glut-RA versus Glut-Arco-1, Glut-HVC-1 versus Glut-NC-1, and Glut-HVC-2 versus Glut-NC-4, for a total of 98,593 DARs (FDR < 0.01, log2 fold-change > 0.6) (**Fig. 3c,d**). The number of DARs between song and non-song neurons exceeded that between pairwise combinations of non-song glutamatergic neurons (**Extended Data Fig. 4c**), indicating that the gene regulatory states of song neurons are strongly differentiated from that of their non-song sister cell types.

We examined the enrichment of transcription factor binding site (TFBS) motifs across the DARs for each song neuron and its paired non-song type and found striking similarities across cell types (**Fig. 3e,f** and **Extended Data Fig. 4d,e**). We found that each comparison yielded similar sets of TFBS motifs that had either higher or lower accessibility in song neurons relative to non-song neurons. Motifs enriched in song neurons included those associated with activity-dependent gene expression (FOS, JUN), chromatin remodeling (SMARCC1), steroid receptors (AR, NR3C1, PGR), development (MAFs, BACHs, NFIs), and oxidative stress (NFE2, NFE2L2) (**Fig. 3e** and **Extended Data Fig. 4d,e**). Motifs that are enriched in DARs with reduced accessibility in song neurons relative to non-song neurons showed a similarly consistent pattern across the three comparisons. In general, each showed a depletion of several development-associated motifs (NEUROGs, NEURODs, OLIGs, BHLHs, TCFs, TALs, MEF2s, ATOHs, and RFXs) with Glut-HVC-1 showing less pronounced depletion of these motifs in its DARs relative to Glut-RA and Glut-HVC-2 neurons (**Fig. 3e**).

To determine the extent to which these patterns of motif enrichment are shared across each song versus non-song comparison, we performed pairwise correlations of the motif enrichment values for families of transcription factors between each of the song glutamatergic neuron types. We found that motif enrichments are highly correlated across each song neuron type (Pearson’s *r* 0.94-0.98) (**Fig. 3f**), indicating that each type, whether it is an extratelencephalic (Glut-RA), palliopallial (Glut-HVC-1), or palliostriatal neuron (Glut-HVC-2), engages a similar set of gene regulatory programs as part of its specialization. However, the transcription factors predicted to bind to these shared motifs exhibited varying levels of differential expression across song neurons relative to their non-song sister type (**Fig. 3g,h** and **Extended Data Fig. 5a-c**). For example, within the MAF family, Glut-RA expresses MAFB, Glut-HVC-1 expresses MAFF and MAFK, and Glut-HVC-2 expresses MAF. A similar pattern occurs for TCF and BHLH families. Exceptions to this pattern were found with AP-1 associated factors (*FOSL2* shows high expression across cell types), MEF2 family (reduced expression of *MEF2C* across cell types), and androgen receptor (high expression of *AR* across cell types).

### Co-option of gene regulatory networks

The analysis above indicates that song glutamatergic neurons express specialized gene regulatory programs that distinguish them from their non-song neuron counterparts. However, it is unclear what gene regulatory innovations enabled song neurons to acquire these specialized programs. The co-option of existing traits into a new context can drive the evolution of complex traits [25]. GRNs regulate the expression of batteries of genes that have coherent functions, and inducing their activity in a new cellular context can confer new properties to the host cell type.

Given the paired relationships between song and non-song neurons and the distinct gene regulatory profiles of song neurons, we hypothesized that GRN co-option underlies the specialized gene expression profiles and functional properties of song glutamatergic neurons. Two initial predictions follow from this hypothesis. First, we expected to find GRNs that are active in both song glutamatergic neurons (’derived’ cell type) and unrelated cell types (’ancestral’ cell type). Second, we expected that there should be significant overlap between the sets of target genes that a given transcription factor regulates in ancestral and derived cellular contexts. To test this hypothesis, we inferred song system-specific GRNs using SCENIC+, a method that leverages correlations across cells to identify putative regulatory relationships, and used simulated disruptions of gene expression to predict determinants of cell identity [26]. We first performed this analysis on the entire dataset to infer GRNs across all cell types (**Fig. 4a** and **Extended Data Fig. 6a**). GRN activities were similar within cell type classes (glutamatergic and GABAergic neurons, astrocytes, oligodendrocytes, etc.). However, glutamatergic neuron types exhibited varying patterns of GRN activity according to brain region (nidopallium versus arcopallium, song versus non-song).

**Figure 4:**
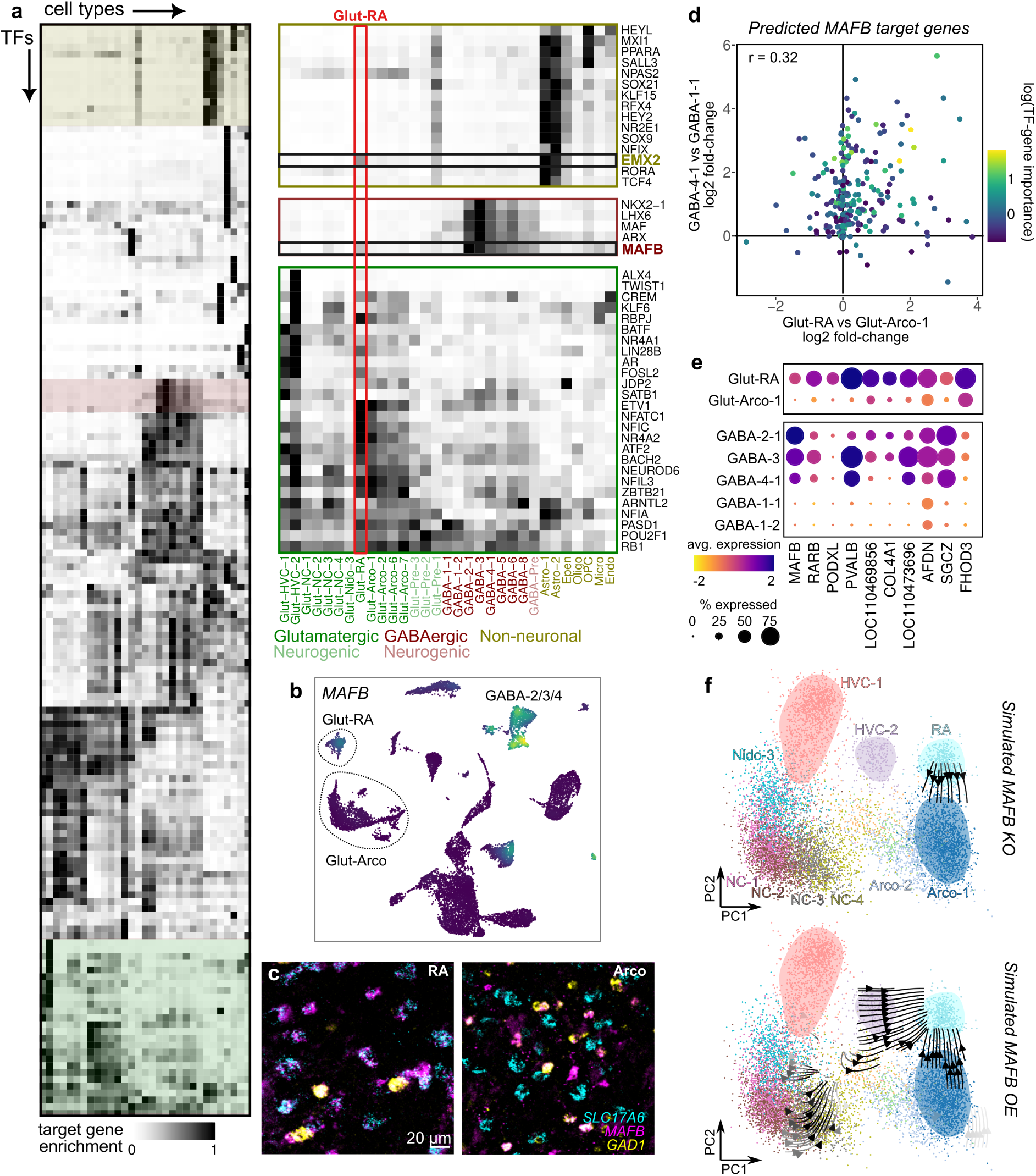
Co-option of GRNs in song motor pathway glutamatergic neurons. **(a)** GRN activity, represented as the enrichment of target gene expression for each transcription factor, in each cell type from the song motor pathway and surrounding non-song motor regions. At right are expansions of the three colored segments of the heatmap representing different patterns of GRN activity. **(b)** UMAP plot of *MAFB* expression. *MAFB* has elevated expression in Glut-RA neurons and in GABA-2/3/4 interneurons (*Sst* and *Pvalb*-class). **(c)** *In situ* hybridization in RA and surrounding Arco for *MAFB*, glutamatergic neurons (*SLC17A6*), and GABAergic neurons (*GAD1*), showing coexpression of *MAFB* in glutamatergic neurons in RA but not in Arco. **(d)** (*Top*) Differential expression (DE) of predicted *MAFB* target genes between Glut-RA and Glut-Arco-1 neurons compared to *Pvalb*-class interneurons (GABA-4-1) and LGE-class interneurons (GABA-1-1). Point color indicates TF to target gene importance from SCENIC+ tree-based regression. **(e)** Expression of selected genes showing elevated expression in Glut-RA and GABA-2/3/4. **(f)** Simulated (*top*) knockout and (*bottom*) overexpression of *MAFB* in glutamatergic neurons. Arrows indicate the direction of shifts in neuron position in PCA space following each manipulation.

We focused on three clusters of GRN activity (**Fig. 4a**). The first cluster exhibited elevated activity across each song glutamatergic neuron relative to non-song glutamatergic neurons. This cluster contains several transcription factors (TFs) identified in the previous analysis, including *AR*, *FOSL2*, and *BATF*. The other two clusters showed strong activity in either GABAergic neurons or astrocytes. In these clusters, two transcription factors, *MAFB* and *EMX2*, showed unexpected activity in Glut-RA neurons. *MAFB* itself is strongly expressed in the GABA-2/3/4 interneuron clusters, homologous cell types to mammalian *Sst-* and *Pvalb*-class neurons [20], and in Glut-RA neurons but not in any other glutamatergic neuron type (**Fig. 4b,c** and **Extended Data Fig. 6b**). Similarly, *EMX2* is strongly expressed in astrocytes and exclusively in Glut-RA neurons relative to other glutamatergic neuron types (**Extended Data Fig. 6c,d**).

We hypothesized that Glut-RA neurons co-opt *MAFB* and *EMX2* GRNs as part of their specialization. If this hypothesis is true, we expect to find that similar genes are targeted by each transcription factor in its “ancestral” context (GABAergic neurons or astrocytes) and its “derived” context (Glut-RA neurons). Indeed, we found that the predicted target genes for *MAFB* exhibited elevated expression in Glut-RA and GABA-4-1 neurons relative to comparator cell types, and those for *EMX2* exhibited similarly elevated expression in Glut-RA and Astro-1 (**Fig. 4d,e** and **Extended Data Fig. 6e,f**). To predict whether the loss of MAFB or EMX2 expression affects the transcriptional specialization of Glut-RA neurons, we used SCENIC+ to simulate knocking out or overexpressing each gene and then assessed how these manipulations affected predicted target gene expression. Simulated knockouts of *MAFB* or *EMX2* shifts the Glut-RA expression profile toward that of Glut-Arco-1, reflecting a de-specialization of song neuron transcriptional identity (**Fig. 4f** and **Extended Data Fig. 6g**). Conversely, overexpressing *MAFB* or *EMX2* pushes glutamatergic neurons toward the specialized Glut-RA state.

Together, these results support a model in which Glut-RA neurons engage conserved gene regulatory networks for *MAFB* and *EMX2* that are normally expressed in other cellular states and that drive the specialization of song neurons from the ancestral non-song state.

### Overexpression of *MAFB* and *EMX2* in chicken glutamatergic neurons induces song neuron expression profiles

To test whether *MAFB* or *EMX2* is sufficient to induce the expression of genes with elevated expression in Glut-RA neurons, we overexpressed each transcription factor (each of the Bengalese finch sequence) in the telencephalons of chicken embryos using a doxycycline-inducible *piggyBac* plasmid (**Fig. 5a** and **Extended Data Fig. 7a-c**) [27]. We injected and electroporated each construct into the telencephalic lateral ventricle of embryonic day 4 (E4) embryos then induced expression using doxycycline at E6. The following day, we microdissected fluorescent tissue and performed single-nucleus RNA-seq. We detected *MAFB* and *EMX2* transgene expression primarily in progenitor cells and glutamatergic and GABAergic neurons (**Fig. 5b** and **Extended Data Fig. 7d**), consistent with stable integration of the plasmids into neural progenitors and their progeny. Within chicken glutamatergic neurons, MAFB or EMX2 over-expression increased the expression of genes predicted to be regulated by MAFB or EMX2, respectively, in songbird Glut-RA neurons, significantly more so than other genes in other predicted regulatory networks (**Fig. 5c**, MAFB: p = 4e-4, EMX2: p = 6e-3, two-sided Komolgorov-Smirnov test). Furthermore, there was a significant overlap between MAFB- and EMX2-induced genes in chicken glutamatergic neurons and those differentially expressed between Glut-RA and its sister non-song neuron type Glut-Arco -1 (**Fig. 5d**). The genes with the strongest induction by each transgene exhibited enriched expression in Glut-RA versus arcopallial neuron types as well as, in several cases, elevated expression in HVC projection neurons relative to their sister cell types (**Fig. 5e,f**). These results indicate that despite the evolutionary distance between chickens and songbirds and the absence of a vocal learning circuit in chickens, the heterologous expression of *MAFB* and *EMX2* in chicken glutamatergic neurons is sufficient to partially drive a song neuron-like expression profile.

**Figure 5:**
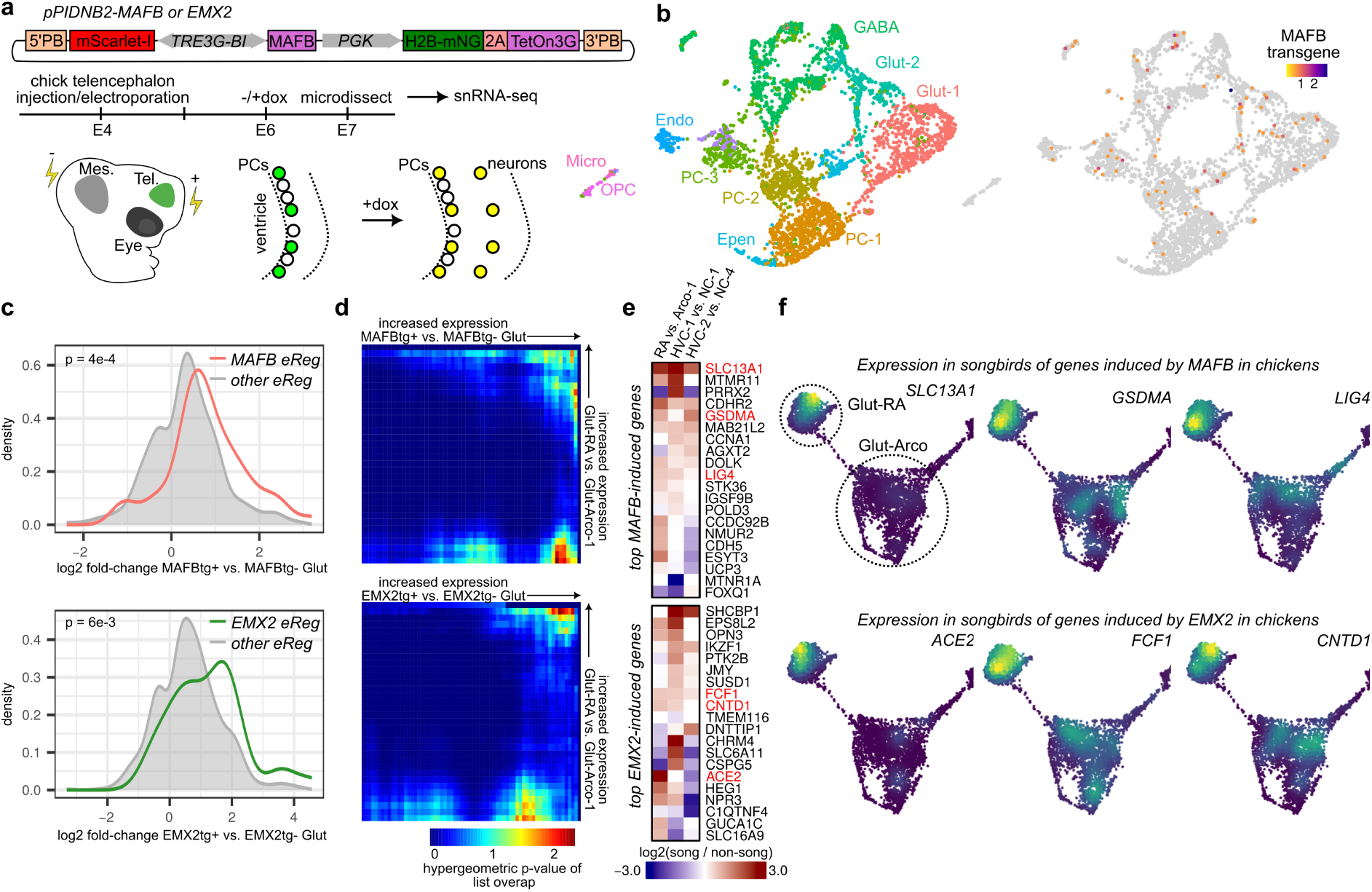
Overexpression of MAFB or EMX2 in glutamatergic neurons induces Glut-RA expression program. **(a)** Experiment design and timeline. A doxycycline-inducible *piggyBac* construct was electroporated into the chicken telencephalon to drive MAFB expression followed by snRNA-seq to assess the transcriptional influence of MAFB in glutamatergic neurons. Tel: telencephalon, Mes: mesencephalon. **(b)** UMAP plots showing (*left*) cell clusters and (*right*) the expression of the MAFB transgene **(c)** Induction of the expression of finch MAFB or EMX2 eRegulon genes by MAFB or EMX2 overexpression in chicken glutamatergic neurons compared to that of genes in other finch eRegulons. **(d)** Rank-rank hypergeometric overlap (RRHO) analysis of differential expression between Glut-RA and Glut-Arco-1 in finch versus (*top*) MAFBtg+ versus MAFBtg- or (*bottom*) EMX2tg+ vs. EMX2tg- glutamatergic neurons in chicken. Higher values in the upper right corner indicate significant overlap of the top differentially expressed genes in each list. **(e)** Differential expression between song and non-song glutamatergic neurons of the top 20 transgene-induced genes. Gene names in red are those depicted in panel **f**. **(f)** UMAP plots of finch RA and arcopallial glutamatergic neurons (as in Fig. 1d) showing the expression of selected transgene-induced genes.

### Song neuron-specific regulation of MAFB and *EMX2*

GRN co-option could occur through the evolution of *cis*-regulatory elements (CREs) that enable transcription factor expression in a new cellular context. If *MAFB* and *EMX2* acquired CREs to drive expression in Glut-RA neurons, we expected to find distinct patterns of chromatin accessibility linked to each gene locus (**Fig. 6a**). We first examined the predicted inputs to each transcription factor in Glut-RA projection neurons, GABA-4 interneurons, and astrocytes using SCENIC+ analyses focused on these cell types. We found that *MAFB* and *EMX2* are predicted to be regulated by distinct sets of TFs in Glut-RA neurons compared to interneurons or astrocytes (**Fig. 6b**). Notably, one of the predicted regulators for MAFB expression in Glut-RA neurons is androgen receptor (*AR*), which is elevated in song neurons and plays a key role in song system maturation and birdsong development [28, 29].

**Figure 6:**
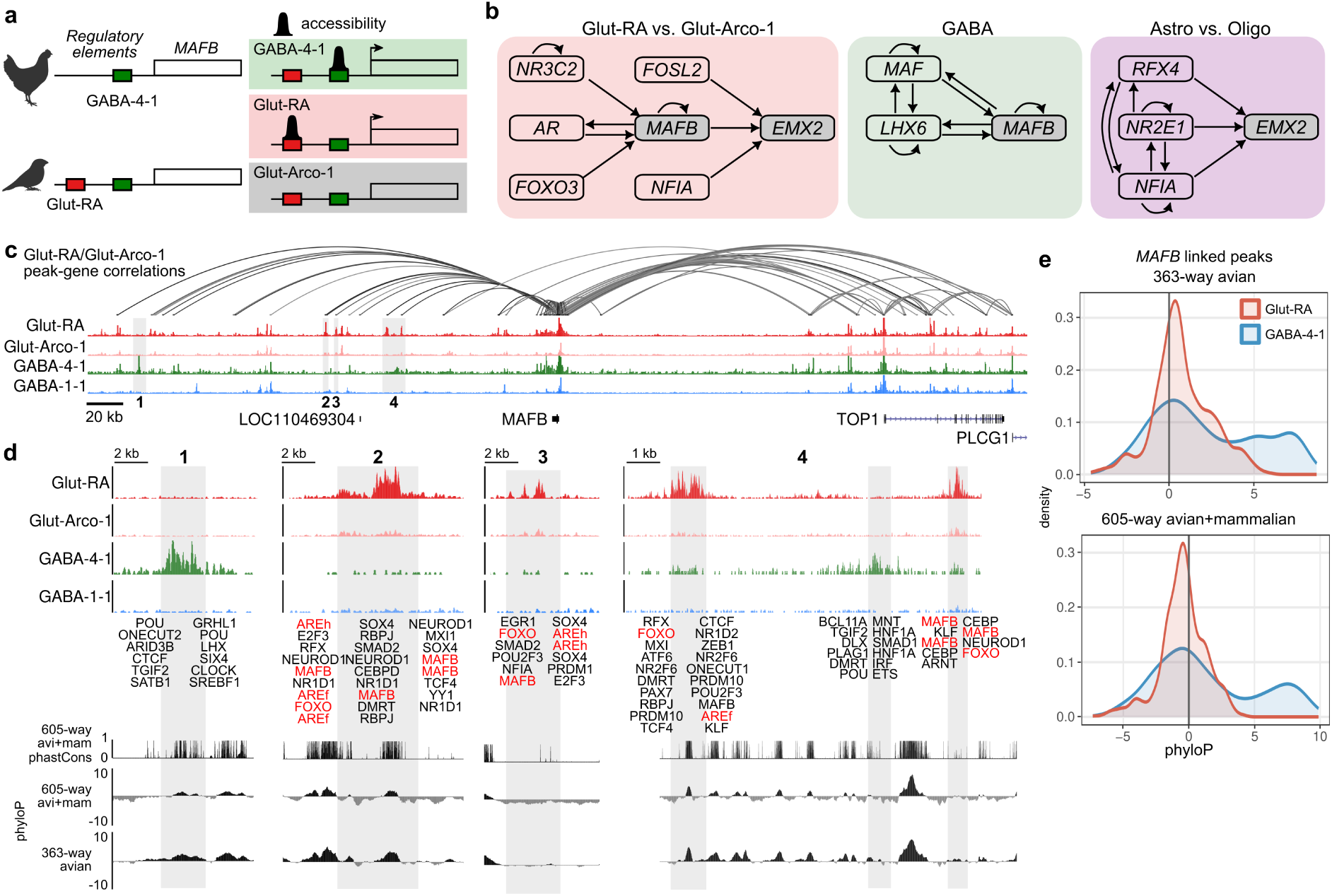
Song neuron-specific regulatory landscape of *MAFB*(a) Model of gene regulatory network co-option through cis-regulatory evolution. (*Left*) Birds without complex vocalizations (e.g. chickens) possess cis-regulatory elements for a putative fast-spiking regulator active in GABA-4-1). However, songbirds have gained a separate set of regulatory elements that can be accessed within Glut-RA neurons. (*Right*) Predicted patterns of chromatin accessibility in songbird neuronal types. **(b)** Predicted SCENIC+ regulatory inputs to *MAFB* and *EMX2* in Glut-RA neurons, GABA-4-1 neurons, and astrocytes. **(c)** Chromatin accessibility flanking the MAFB locus. Arcs above the accessibility tracks link the MAFB transcription start site with DARs whose accessibilities covary with *MAFB* expression (Pearson r > 0.4). **(d)** Magnification of the numbered regions indicated in panel **b**. Transcription factor binding sites for each differentially accessible region are annotated. Sites for SCENIC+ predicted inputs are highlighted in red. AREh, androgen receptor element half-site; AREf, androgen receptor element full-site. At bottom are three tracks showing the conservation (phastCons) or conservation and acceleration (phyloP) across two alignments, one containing 605 species of birds and mammals (avi+mam) and a second with 363 species of birds only. **(e)** Distributions of phyloP scores across *MAFB*-linked DARs with high differential accessibility (normalized fragment density differences greater than 5 between Glut-RA and Glut-Arco-1 or GABA-4-1 and GABA-1-1).

Next, we examined the chromatin accessibility profiles around each gene locus between Glut-RA neurons and the respective non-song cell types for each factor, fast-spiking interneurons (GABA-4-1) and astrocytes (Astro-1) (**Fig. 6c,d** and **Extended Data Fig. 8a,b**), relative to tracks for comparator cell types for each principal type: Glut-Arco-1, an LGE-class interneuron (GABA-1-1), and oligodendrocytes (Oligo). Consistent with each gene having distinct regulatory inputs in derived and ancestral cell types, we found that each locus contains CREs that are solely active in Glut-RA neurons and not in their paired non-song neuron cell type, Glut-Arco-1. Furthermore, for each locus, a subset of these CREs is accessible in Glut-RA neurons, yet inaccessible in GABA-4 or Astro-1 cell types.

If *MAFB* and *EMX2* are activated by recently evolved regulatory elements in Glut-RA neurons, we expected to find differential rates of conservation in Glut-RA DARs relative to DARs specific for GABA-4-1 interneurons or astrocytes. To assess this prediction, we analyzed conservation signals (both phastCons and phyloP) across two whole-genome alignments: one containing 363 assemblies from the B10K project that span the avian phylogenetic tree [30] and a second that combines this avian alignment with a 242-species mammalian alignment from the Zoonomia consortium [31], yielding 605 species (**Fig. 6c**). Glut-RA DARs that were linked to *MAFB* showed weak conservation across avians and mammals (**Fig. 6e**). In contrast, regions with high accessibility in GABA-4-1 interneurons relative to GABA-1-1 interneurons tended to be deeply conserved across birds and mammals. *EMX2*-linked DARs showed less pronounced differences in conservation signatures between groups, yet astrocyte-specific DARs showed slightly higher conservation across birds and mammals relative to Glut-RA-specific DARs (**Extended Data Fig. 8c,d**). Combined, these results indicate that the *cis*-regulatory inputs to *MAFB* and *EMX2* in song-dedicated projection neurons are distinct from and less conserved than those in fast-spiking interneurons and astrocytes, respectively.

## Discussion

Specialized sensorimotor circuits are necessary for the learning and performance of a range of complex motor skills. However, the molecular factors that drive their divergence from other sensorimotor circuits are poorly understood. Here, we show that glutamatergic neurons differ substantially between brain regions dedicated to birdsong and adjacent regions associated with other sensorimotor behaviors. Each of these specialized song neuron types has a base transcriptional profile that is similar to a non-song neuron type, suggesting that song and non-song neurons represent paired sister cell types that are a product of cell type evolution. Comparisons with neurons in the chicken brain indicated that non-song neurons in songbirds are transcriptionally similar to glutamatergic neurons in chickens, supporting a model in which song neurons are derived cell types specific to songbirds.

The diversity of cell types allows for functional specialization. However, it is largely unclear how new cell types evolve. Cell types are proposed to evolve in a stepwise manner through diversification into sister cell types, each with regulatory independence and subject to their own functional divergence [24, 32]. Recent evidence enabled by advances in single-cell sequencing technologies suggests that differences in the cell type composition of brains and other organs contribute to the evolution of behavior, such as in the cases of the Mexican tetra (*Astyanax mexicanus*), *Drosophila yakuba*, and oldfield mouse (*Peromyscus polionotus*) [33–36]. In songbirds, the sister cell type organization of song and non-song glutamatergic neurons and the dedication of song neurons to a clade-specific behavior provides an avenue to understand the mechanisms underlying cell type diversification and its role in behavioral evolution. We found that song glutamatergic neurons express a shared set of gene regulatory programs that differentiate them from their non-song comparators. The identification of this shared set of gene regulatory programs touches on an open question as to how such a large circuit comprising multiple regions could evolve in a coordinated manner. One popular hypothesis suggests that the song system evolved through the duplication and divergence of an ancestral motor circuit that coordinates avian movement [6, 7], as supported by the observation of circuit-level duplication throughout neurobiological evolution [37, 38]. Our data suggests that song neuron specification occurs through the induction of a shared set of transcription factor families across the various song glutamatergic neuron types. Within this common regulatory framework, the differential expression of paralogs from these shared families drives the functional specialization of each cell type within the circuit. This pattern is in line with previous work that demonstrated that singing drives a shared gene regulatory response across each song region that is followed by an induction of region-specific transcription factors [39].

We identified two cases of this cell-type specificity in the expression of MAFB and EMX2 in RA projection neurons (Glut-RA), extratelencephalic projection neurons homologous to mammalian layer 5 subcerebral projection neurons [20]. Glut-RA neurons project to brain-stem regions controlling syringeal and respiratory motor neurons, and their activity is correlated with the acoustic output of song [40, 41]. Glut-RA neurons have fast-spiking electrophysiological properties, including narrow action potential widths and resurgent sodium currents, mediated by Kv3 and Navβ4 channels, respectively, which are believed to enable the high spectrotemporal precision of birdsong [18, 42]. Notably, these physiological properties are characteristic of mammalian Betz cells, large subcortical projection neurons that are abundant in the primate motor cortex but sparse in rodent brains [43] and may support fine motor control [1]. Furthermore, these properties are hallmarks of fast-spiking parvalbumin-positive (PV) GABAergic interneurons [44]. The loss of MAFB disrupts the differentiation of PV cortical interneurons and increases cortical excitability in mice [45, 46]. Premature MAFB overexpression in embryonic mouse spiral ganglion neurons – another fast-spiking cell type – accelerates clustering of AMPA receptors at synaptic terminals [47], which is critical for rapid glutamatergic transmission that supports high firing rates. These converging roles in both fast-spiking interneuron specification and AMPA receptor trafficking make the high expression of MAFB that we observe in fast-spiking song projection neurons particularly striking, suggesting that MAFB may be a conserved regulator of fast-spiking identity across circuits.

How might these regulatory programs be deployed in Glut-RA neurons? Co-option – the reuse of existing gene-regulatory programs in novel contexts – provides a plausible mechanistic explanation through changes to the regulatory environment surrounding an ancestral locus [48, 49]. For example, the acquisition of accessible enhancer regions around the MAFB locus in Glut-RA neurons could confer cell-type-specific expression of this regulatory factor and its targets, giving rise to the unique molecular and physiological signature of this neuron population. Characterization of this locus in Glut-RA neurons suggests that the acquisition of androgen receptor (AR) binding sites (**Fig. 6**) may represent this type of regulatory innovation, driving the high MAFB expression that defines this cell type. AR has long been recognized as an important factor in the development of the male song circuitry in sexually dimorphic songbird species [29, 50, 51]. Taken together, this suggests that the acquisition of AR-mediated regulation at the MAFB locus may act as a mechanism for the co-option of this regulatory network, a key step in establishing the molecular identity of Glut-RA neurons.

The partial induction of song neuron expression profiles in embryonic chicken telencephalon following MAFB overexpression further supports a co-option model, suggesting the evolution of the song circuit involved the recruitment of regulatory components available in a common ancestor rather than their *de novo* invention. A similar argument can be made for the independent evolution of vocal learning in humans. Non-human primates largely produce innate vocalizations which lack the flexible, learned character of human speech [52]. Crucially, the difference in the location of the laryngeal motor cortex (LMC) in humans and non-human primates is hypothesized to have enabled the establishment of a direct connection between the LMC and brainstem motoneurons necessary for rapid coordination between laryngeal, orofacial, and respiratory movements underlying speech production [53]. This organization bears striking resemblance to the projection of Glut-RA neurons to brainstem regions controlling syringeal and respiratory motor neurons [10], which is notable given the identification of RA and LMC as functional and molecular analogs in vocal-learners [19]. The shared presence of these specialized characteristics that enable the production of vocalizations across species raises the possibility that similar genetic innovations may have contributed to the evolution of human speech circuitry.

## Methods

### Animal use

All Bengalese finches were from breeding colonies at UCSF or UCSC, or were purchased from approved vendors. Experiments were conducted in accordance with NIH, UCSF, and UCSC policies governing animal use and welfare.

### Single-nucleus RNA-sequencing (snRNA-seq)

Three adult (>120 days post hatch) male Bengalese finches were euthanized via isoflurane overdose then decapitated. The brain was dissected out into ice-cold ACSF (152 mM choline chloride, 2.5 mM KCl, 1.1 mM NaH_2_PO_4_, 25 mM NaCO_3_, 35 mM glucose, 10 mM HEPES, 3 mM MgSO_4_*7H_2_O, and 0.5 mM CaCl_2_*2H_2_O) bubbled with carbogen. The brain was then sectioned in ice-cold carbogenated ACSF either coronally or sagittally using a Leica VT1000 vibratome at 300 *µ*m. Areas of interest were microdissected out, placed in cryovials, frozen in liquid nitrogen, then transferred to −80*^◦^*C for long-term storage. Nuclei were isolated from microdissected tissue using the method described in [20]. 500 *µ*L ice-cold lysis buffer (10 mM Trizma pH 7.4 (Sigma, T2194), 10 mM NaCl, 3 mM MgCl_2_, and 0.1% NP-40) was added to each cryovial, and the tissue was pooled across individual birds into 2 mL glass dounces (Kimble). Samples were dounced 15 times using a loose pestle then 15 times using a tight pestle. 900 *µ*L sucrose cushion buffer (Sigma, NUC201-1KT) was added to each sample and was slowly mixed by pipetting 20 times. 500 *µ*L sucrose cushion buffer was added to a new 2 mL tube, then the 1400 *µ*L of tissue/lysis/sucrose bufffer was carefully layered on top. Samples were centrifuged for 45 minutes at 13,000xg at 4*^◦^*C. All but 100 *µ*L supernatant was discarded, 300 *µ*L nuclei suspension buffer (1x PBS, 1% UltraPure BSA (ThermoFisher, AM2616), and 0.5% RNase inhibitor (Clontech, 2313A)) was added, then the pellet was gently resuspended by pipetting. Samples were centrifuged for 5 minutes at 550xg at 4*^◦^*C. All but 70 *µ*L of supernatant was discarded, the pellet was resuspended, and the sample was passed through a 40 *µ*m FlowMi filter (Bel-Air). The nuclei were counted on a C-Chip and standardized to 400-700 nuclei/*µ*L. Nuclei were added to target a recovery of 10,000 nuclei onto a Next GEM Chip G microfluidics chip as part of the Next GEM Single Cell 3’ Kit v3.1 (10X Genomics). Following multiplexing, libraries were sequenced on two lanes of an Illumina NovaSeq S4 200 flow cell (101 cycles R1, 12 cycles I1, 24 cycles I2, 101 cycles R2) at the UCSF Center for Advanced Technology.

### Single-nucleus RNA and ATAC-sequencing (Multiome-seq)

For snRNA/ATAC-seq samples, seven adult (>120 dph) male Bengalese finches were anesthetized with isoflurane, transcardially perfused with ice-cold carbogenated ACSF, then decapitated. Brains were dissected out into ice-cold carbogenated ACSF then coronally sectioned using a Leica VT1000 vibratome at 300 *µ*m. Areas of interest were microdissected out, placed in cryovials, frozen in liquid nitrogen, then transferred to −80*^◦^*C for long-term storage. Nuclei were isolated using a modified version of the Optiprep method [54, 55]. Several stable buffers were prepared before nuclei isolation and stored a 4*^◦^*C: 1.079x Homogenization Buffer Stable Solution (”HBS”, 260 mM sucrose, 30 mM KCl, 10 mM MgCl_2_, 20 mM Tricine-KOH pH 7.8), Diluent Buffer (150 mM KCl, 30 mM MgCl_2_, 120 mM Tricine-KOH pH 7.8), 50% iodixanol solution (diluted from 60% Optiprep Density Gradient Medium, Sigma, D1556, using Diluent Buffer), and 5X ATAC-RSB (50 mM Tris-HCl pH 7.5, 50 mM NaCl, 15 mM MgCl_2_). On the day of nuclei isolation, the following buffers were prepared: 1x Homogenization Buffer Unstable Solution (”HBU”, HBS to 1x, 1 mM DTT, 0.5 mM spermidine, 0.15 mM spermine, 0.3% NP-40, 1 U/*µ*L Protector RNase Inhibitor (Sigma 3335402001), 1x Complete Protease Inhibitor (prepared from a 25x solution consisting of one tablet (Sigma, 11697498001) dissolved in 2 mL nuclease-free water)), 30% Iodixanol Solution and 40% Iodixanol Solution (both diluted from 50% iodixonal solution using 1x Homogenization Buffer Unstable Solution), and ATAC-RSB-Tween (1x ATAC-RSB, 0.1% Tween-20, 1% BSA, and 1 U/*µ*L Protector RNase Inhibitor). Samples from different birds were pooled and 1 mL 1x HBU was added. Samples were incubated on ice for 5 minutes, then transferred to a 2 mL glass dounce (Kimble). Samples were homogenized with 15 strokes of the loose pestle, then 15 strokes of the tight pestle. Samples were transferred to 2 mL tubes then centrifuged for 5 minutes at 350xg at 4*^◦^*C using a swing-out rotor. All but 200 *µ*L of supernatant was removed, the total volume was brought up to 400 *µ*L using 1x HBU, then the pellet was gently resuspended by pipetting. 400 *µ*L 50% iodixanol solution was added and mixed well by pipetting. 600 *µ*L 30% iodixanol solution was layered underneath the 25% iodixonal/tissue layer, followed by 600 *µ*L 40% iodixanol solution underneath the 30% layer. In a pre-chilled swing-out rotor, samples were centrifuged for 20 minutes at 10,000xg at 4*^◦^*C with the brake off. Starting at the top of the solution, enough solution was aspirated off to leave 200-300 *µ*L above the interface between the 30% and 40% iodixanol layers, which contains nuclei. 200 *µ*L of the nuclei band was aspirated and transferred to a clean 1.5 mL tube containing 1 mL of ATAC-RSB-Tween. Nuclei were centrifuged for 10 minutes at 500xg at 4*^◦^*C using a swing-out rotor. The entire supernatant was carefully removed, followed by addition of 12 *µ*L Nuclei Resuspension Buffer (10X Genomics) and gentle resuspension by pipetting.

Nuclei were counted and their concentrations were adjusted to 800-12,000 nuclei/*µ*L. The nuclei were then loaded onto a Next GEM Chip J (10X Genomics) at numbers targeting 2,500 to 10,000 recovered nuclei. Sequencing libraries for each region were prepared using the Chromium Next GEM Single Cell Multiome ATAC + Gene Expression kit. The libraries were then pooled, and sequenced on an Illlumina NovaSeq S4 200 flow cell at the UCSF CAT.

### snRNA-seq preprocessing

FASTQ files were aligned to the Bengalese finch reference genome (lonStrDom2, GCF_005870125.1) using *cellranger v7.1.0*. Included also in the alignment index were gene models with extended 3’ UTRs as described in [56]. Ambient RNA was removed using *cellbender v0.3.2* [57] with a false positive rate of 0.01. Using *Seurat v5.2.1*, nuclei were removed that had greater than 2% mitochondrial reads, greater than 20,000 UMIs, and less than 1000 UMIs. Doublets were removed using two methods. First, nuclei were demultiplexed by their genotype using *souporcell v1.0* [58], which assigns each nucleus to an individual bird and estimates whether a nucleus is a doublet or singlet. Second, we performed doublet detection using *DoubletFinder v2.0.4* [59]. Nuclei were retained if they were called a singlet in both approaches. Gene expression was then normalized using *SCTransform*, transformed via principal components analysis, then reduced using uniform manifold approximate projection (UMAP) and the first 30 PCs. Clustering was performed iteratively on progressively smaller subgroups to identify subclusters. First, clustering was performed on the entire dataset using Louvain clustering with multilevel refinement and a resolution of 0.2 using the Seurat function *FindClusters*. At this level, major cell classes were identified based on marker gene expression (e.g. glutamatergic and GABAergic neurons, astrocytes, oligodendrocytes, etc.). Glutamatergic neurons were subsetted and reprocessed via *SCTransform*, PCA, UMAP, and clustering to identify clusters belonging to regional domains, nidopallium and arcopallium. Glutamatergic neurons in each region were then similarly reprocessed to identify region-specific subclusters. GABAergic neurons were not subdivided by region but were similarly subsetted and reprocessed as the glutamatergic neurons.

### Cross species integration

Single-nucleus RNA-sequencing data from the adult chicken telencephalon was downloaded from https://heidata.uni-heidelberg.de/dataset.xhtml?persistentId=doi:10.11588/DATA/BX6REK [23]. We filtered the chicken dataset to include only glutamatergic (”Ex”) clusters while also omitting “Ex_TCF7L2”, which was thought to be a thalamic population in the original publication. Likewise, we filtered the finch dataset to only include glutamatergic (”Glut”) neurons. We then filtered each dataset for one-to-orthologs between chicken and finch. One-to-one orthologs between chicken (*Gallus gallus*) and zebra finch (*Taeniopygia guttata*) were downloaded using the Genome Pair Viewer utility on the OMA Browser (https://omabrowser.org/oma/genomePW/, [60]) accessed on April 8, 2025. Zebra finch is an estrildid finch and closely related to the Bengalese finch. We transferred labels using *Seurat v5.2.1* function *FindLabelTransfers* (normalization.method = “SCTransform”, k.anchor = 10, k.filter = 50, k.score = 30, max.features = 200, reduction = “rpca”, dims = 1:40) followed by *TransferData*. We then averaged the transfer probabilities within each pair of chicken and finch clusters to obtain an aggregate label transfer value. The maximum of these values was then found for each finch cluster to determine their overall degree of similarity to chicken glutamatergic neurons as a whole.

### Multiome-seq preprocessing

FASTQ files were aligned to the Bengalese finch reference genome (lonStrDom2, GCF_005870125.1) using *cellranger-arc v2.0.2* and the gene model GTF with extended 3’ UTRs described above. Ambient RNA was removed using *cellbender v0.3.2* [57] with a false positive rate of 0.05. Using *Seurat v5.2.1*, nuclei were retained that satisfied the following conditions: 1000 < number of GEX UMIs < 25,000, 1000 < number of ATAC fragments < 30,000, nucleosome signal < 2, and TSS enrichment >1. Doublets were removed using the approach in *snRNA-seq pre-processing*. *Souporcell* and *DoubletFinder* were used to independently identify probable doublets, and we only retained nuclei that were labeled singlet by each method. Cluster labels were assigned using Seurat label transfer of labels from the snRNA-seq dataset.

### Regional cell type specificity

The regional specificity (as in Figure 1f) for each cell type T was calculated as Σ(*x_RT_* ∗*log*(*x_RT_* /*n*)). The value *x_RT_* was obtained by first normalizing the number of cells of type T by the total number of cells in a region R, yielding the proportion of cell type T in region R. For a given cell type T, these values were normalized across regions by their sum to give *x_RT_* (max value 1). *n* is the number of regions.

### Differential gene expression analysis

To identify differentially expressed genes between clusters, we first generated pseudobulk clusters using the Seurat function *AggregateExpression* with nuclei grouped by cluster and biological replicates (3 birds for snRNA-seq data and 7 birds for Multiome-seq data). The assignment of each nucleus to a biological replicate was obtained using *souporcell* as described in *snRNA-seq preprocessing*. Differentially expressed genes across clusters were then found using the Seurat function *FindMarkers* with the *DESeq2* method [61]. For the hierarchical clustering of glutamatergic neurons, we first identified differentially expressed genes for each Glut neuron type relative to all other Glut neuron types. We then removed genes that were differentially expressed (adjusted p-value < 0.1, Benjamini-Hochberg correction) in song neurons (Glut-HVC-1, Glut-HVC-2, Glut-LMAN-1, Glut-LMAN-2, or Glut-RA). Then for each cluster, we selected genes that had an adjusted p-value < 0.1 and had an absolute average log2 fold-change average that was within either the top 100 overexpressed or underexpressed genes for that cluster. We hierarchically clustered the resulting matrix using the *R* function *pvclust* (method.dist = “correlation”, method.hclust = “ward.D2”, nboot = 1000) [62].

For the analysis of the effects of MAFB or EMX2 overexpression, we used *FindMarkers* with the Wilcoxon Rank Sum test comparing transgene-positive versus transgene-negative glutamatergic neurons. To determine the significance of gene overlap between lists of differentially expressed genes, we used rank-rank hypergeometric overlap (RRHO) using the R package *RRHO2 v1.0* [63]. Genes were ordered by their log2 fold-change between conditions.

### Accessible region analysis

snATAC-seq analysis was performed using *ArchR v.1.0.3* [64]. CRE regulation of nearby genes was predicted using the *ArchR* function *getPeak2GeneLinks* (corCutOff = 0.4).

### Differential accessibility

To identify accessible regions, we first generated pseudobulk replicates for each cluster using the function *addGroupCoverages* then using *macs2 v2.2.9.1* via the *ArchR* function *addReproduciblePeakSet*. To identify differentially accessible regions, we used the function *getMarkerFeatures* (Wilcoxon test, bias = c(”TSSEnrichment”, “log10(nFrags)”)) and filtered for regions with an FDR <= 0.1.

### Motif enrichments

To identify the enrichment of transcription factor binding site (TFBS) motifs across accessible peaks, we first annotated each peaks with potential binding sites using the *ArchR* function *addMotifAnnotations* and the CIS-BP database for *Homo sapiens* [65]. Then, we used *peakAnnoEnrichment* (which performs hypergeometric tests comparing motif presence within a subset of peaks relative to all peaks in a set) to identify motifs that are enriched in peaks that were either more accessible (FDR <= 0.01 and log2 fold-change >= 0.6) or less accessible (FDR <= 0.01 and log2 fold-change <= −0.6) in song glutamatergic neurons (Glut-HVC-1, Glut-HVC-2, and Glut-RA) relative to their non-song pairs (Glut-NC-1, Glut-NC-4, and Glut-Arco-1). To determine the relative enrichment of TFBS motifs in each glutamatergic cluster, we performed *chromVAR v1.28.0* [66] using the *ArchR* function *addDeviationsMatrix*, which calculates the deviations of each motif present in the accessible regions of a given cell relative to a set of background regions pulled from all cells that are matched by GC-content and accessibility.

### Accessibility tracks

UCSC Genome Browser chromatin accessibility bigwig tracks were generated using *Snap-ATAC v2.8.0* [67] using insertion sites, a bin size of 10, and a smoothing window of 20.

### Gene regulatory network inference

Gene regulatory networks were predicted using *SCENIC+ 1.0a1* [26]. We first analyzed Multiome-seq data using *pyCisTopic 2.0a0* (1) to call pseudobulk peaks via *macs2*, (2) to identify sets of differentially accessible region, and (3) to identify topics of variable accessibility across cells. When then generated a custom cisTarget database of motif rankings and scores using *cisTarget* with accessibility peaks identified across all cells in the dataset. SCENIC+ was run using default parameters except for the following: motif_similarity_fdr = 0.01 (default 0.001) and ctx_rank_threshold = 0.2 (default 0.05). This pipeline was used to infer GRNs for subsets of cell types: all glutamatergic neurons, Glut-RA vs. Glut-Arco-1 neurons, GABAergic cells, and between astrocytes and oligodendrocytes.

For *in silico* perturbations of gene expression, we used SCENIC+ function to train TF regressors for the top 5000 genes according to triplet rank. For simulated knockdown, the expression of either *MAFB* or *EMX2* was set to 0; for overexpression, expression was set to 10. Simulated perturbations were calculated using *simulate_perturbation* for five iterations. Predicted perturbations in PCA space were computed using the function *plot_perturbation_effect_in_embedding*.

### *In situ* hybridization

Fluorescent *in situ* hybridization via hairpin chain reaction (FISH-HCR) was performed using cryosectioned adult (>120 days old) Bengalese Finch brain tissue. Adult Bengalese finches were euthanized via isoflurane overdose then decapitated. Brains were dissected out, flash frozen in isopentane at −70°C for 12 seconds within 5 minutes of decapitation, and stored at −80°C. Frozen brains were cryosectioned into 16 µm thin sections onto Superfrost Plus slides (Fisher), which were then stored at −80°C. HCR probes, buffers, and hairpins were purchased from Molecular Instruments, except Probe Wash Buffer (PWB). PWB contains 50% formamide, 5X SSC, 9% 100mM Citric Acid (pH 6.0), 0.001% Tween 20, 0.5% Heparin (10mg/mL initial concentration), and 15% dH2O. Cryosectioned tissue was fixed on microscope slides using ice cold 4% paraformaldehyde (PFA)/phosphate-buffered saline (PBS) for 15 minutes. The tissue was then washed three times for 5 minutes in PBS + 0.1% Tween-20 (PBST) on a shaker. The tissue was then dehydrated using a series of ethanol (EtOH) washes on a shaker at room temperature (RT): one 3 minute wash of 50% EtOH, one 3 minute wash of 70% EtOH, and two 3 minute washes of 100% EtOH. The tissue was allowed to air dry for 3 minutes at RT. Next, the tissue was incubated at 37°C with 150 µL probe hybridization buffer for 10 minutes, then incubated overnight at 37*^◦^*C in 150 µL probe hybridization buffer containing 4 nM of each probe. The next day, tissue was submerged in PWB for 5 minutes at RT to release coverslips from the tissue. Next, the tissue was washed through a series of PWB/SSCT (5× sodium chloride sodium citrate + 0.1% Tween 20) for 15 minutes per wash at 37°C in the following order: 75%PWB/25%SSCT, 50%PWB/50%SSCT, 25%PWB/75%SSCT, 100% SSCT. The tissue was then washed once more with SSCT for 5 minutes at RT. 200 µL amplification buffer was applied, and the tissue was incubated at RT for 30 minutes. Fluorophores were incubated at 95°C for 90 seconds and left to rest in the dark for 30 minutes at RT before being applied to the tissue in 110 µL amplification buffer. The tissue was incubated in the dark at RT overnight. Finally, on the third day, the tissue was added to 5× SSCT at RT to remove the coverslips, 200 µL of 5× SSCT + DAPI (1 ug/mL final concentration) was added to then incubate at RT in the dark for 30 minutes. Next, the tissue was washed for 30 minutes in 5× SSCT at RT in the dark before one more wash with 5× SSCT for 5 minutes at RT in the dark. Prolong Glass Antifade Medium (Thermofisher) was applied to each slide and coverslipped. Imaging was performed on a Zeiss 880 Confocal at the UCSC Microscopy Core facility using a 10X objective.

### Cloning

To create pPIDNB2-MAFB and pPIDNB2-EMX2, pPIDBN2 was first linearized using AflII and PstI (NEB). The open reading frames for *MAFB* and *EMX2*, including complementary arms for insertion into pPIDNB2, were synthesized by Twist Biosciences and were then Gibson cloned into pPIDNB2. Correct sequence was verified using whole plasmid sequencing.

### *In ovo* electroporation and snRNA-seq

Fertilized chicken eggs were obtained via overnight shipping and incubated at 37.5 degrees Celsius and 65% humidity for 3 days upon arrival. Eggs were prepped for *in ovo* manipulation following [68] at E4. Injections were performed using a Nanoject III and beveled glass capillaries. The injection solution contained 1 *µ*g/*µ*l of pPIDNB2-MAFB or pPIDNB2-EMX2 over-expression construct, 0.5 *µ*g/*µ*l piggyBac transposase, and 0.1% fast green for visualization in 1x PBS. Embryos were injected with 500 nL of this solution at a rate of 10 nL/s into their telencephala to target radial glial cells. After injection, small holes were punctured through the amnion next to either side of the head using #5 forceps to allow electrode introduction around the embryo. A scoop-shaped positive electrode was positioned underneath the telencephalon and a thin needle-shaped negative electrode was positioned behind and touching the mesencephalon. Embryos were then electroporated with 22 Volts of square wave for 5 pulses at a 50 ms duration with a 0.5 second pulse interval. Eggs were then treated with penicillin-streptomycin to reduce mortality from infection, the egg windows were covered with tape, and the eggs were placed back in the incubator. Successful electroporation was confirmed by GFP signal in the telecephala on E5. Four EMX2 embryos and three MAFB embryos received 200 ul of 100 ng/ml doxycycline on E6 while three EMX2 embryos and two MAFB embryos were left untreated. Following dox-induction, fluorescent telencephalic regions were imaged and microdissected at E7. Fluorescent tissue was flash frozen in liquid nitrogen in 1.5 ml cryovials and stored at −80*^◦^*C for snRNA-seq.

For snRNA-seq, microdissected tissue from chicken embryos was pooled by condition (e.g. tissue from all pPIDNB2-MAFB electroporated, doxycycline treated individuals was combined) before proceeding to nucleus isolation. Nuclei were isolated using a modified version of the Optiprep method [54, 55] as described above in “Single-nucleus RNA and ATAC-sequencing (Multiome-seq)” with an adjusted concentration of 0.2 U/*µ*L Protector RNase Inhibitor and the addition of 10 mg/mL Kollidon VA-64, Sigma, 190845 to HBU. Following isolation, nuclei were washed in 1 mL of Illumina Nuclei Suspension Buffer (NSB) and pelleted by centrifugation at 500 × g for 5 minutes at 4*^◦^*C according to Illumina Single Cell 3′ RNA Prep protocol. Nuclei were resuspended in 20 *µ*L of NSB before counting. Nuclei were counted Revvity Cellometer K2 Fluorescent Cell Counter and diluted to 2500 nuclei/*µ*L in NSB for samples at higher concentrations. We did not recover nuclei for the EMX2/no dox sample and proceeded with the remaining samples.

snRNA-seq libraries were prepared according to the Illumina Single Cell 3′ RNA Prep, T2 protocol. Reactions were overloaded with 10,000 nuclei for samples whose concentrations supported this. Library concentrations were quantified using NEBNext Library Quant Kit for Illumina (NEB, E7630) before pooling. Libraries were sequenced on an AVITI PE150 flow cell at the DNA Technologies and Expression Analysis Cores at the UC Davis Genome Center. FASTQ files were mapped using Illumina DRAGEN 4.4.6000 to a custom chicken reference index consisting of a chicken assembly (bGalGal1.mat.broiler.GRCg7b, GCF_016699485.2) combined with either the pPIDNB2-MAFB or pPIDNB2-EMX2 sequence.

### Western blot

1.6 *µ*g of the pPIDNB2-MAFB or pPIDNB2-EMX2 construct along with 833.33 ng of piggy-Bac transposase were transfected into CFS414 finch fibroblast cells [69] using Lipofectamine 3000 (Invitrogen) into a 6-well cell culture plate. Doxycycline was added the following day at 500 ng/mL. The next day, cells were collected to extract protein using RIPA lysis buffer (0.15M NaCl, 0.03M Tris-HCl, 0.1% NP-40, 12 mM sodium deoxycholate, and 3.5mM SDS) along with 1X CompleteTM protease inhibitor cocktail (Sigma, 11697498001). 12.5 *µ*g protein samples were loaded onto a 10% Criterion XT Bis-Tris Protein Gel along with Precision Plus WesternC Standards ladder (BioRad). The gel was run for 45 min at 200 V and transferred to nitrocellulose paper using a transfer cassette. This was followed by 3x10 minutes of TBS-tween (TBST) washes, blocking with 5% non-fat dry milk in TBST for 1 hour, and blotted using rabbit *α*-MAFB (Millipore, HPA005653-100UL) at 1:1000 dilution or rabbit *α*-EMX2 (ThermoFisher, PA5-103315) at 1:1000 dilution and Beta Tubulin Rabbit PolyAb (Proteintech, 10094-1-AP) at 1:2000 dilution primary antibodies overnight in a 4*^◦^*C shaker. The following day, 3x10 minute washes in TBST were performed and the blot was incubated with goat *α*-rabbit-AP secondary antibody (BioRad, 1706518) at 1:3000 dilution for 2 hours on a room temperature shaker. The blot was developed using Immun-Star AP Chemiluminescence reagents (BioRad, 1705018).

### Immunocytochemistry

166.67 ng of the constructs pPIDNB2-MAFB or pPIDNB2-EMX2 along with the 83.33 ng piggyBac transposase plasmid were transfected into CFS414 finch fibroblast cells on chamber slides using Lipofectamine 3000 (Invitrogen). The following day, 500 ng/mL doxycycline was added to each +dox. The following day, cells were washed 3x10 minutes using DPBS containing calcium and magnesium, fixed using 1% paraformaldehyde (PFA) for 10 minutes, blocked with 5% normal donkey serum-PBS for 1 hour in a room temperature shaker and stained using rabbit *α*-MAFB (Millipore, HPA005653-100UL) at 1:1000 dilution or rabbit *α*-EMX2 (ThermoFisher, PA5-103315) at 1:1000 dilution and mouse *α*-RFP tag monoclonal antibody (Thermo Scientific MA5-15257) at 1:2000 dilution overnight in a 4*^◦^*C shaker. 3x5 minute washes were performed the next day using PBS. Cells were incubated with the respective *α*-rabbit Alexa Fluor™ 647 (Invitrogen A31573) and *α*-mouse Min X Rat CF555 (Biotium 20037) both at 1:2000 dilution secondary antibodies along with DAPI at 1:1000 dilution in a room temperature shaker covered with foil for 1 hour. Cells were washed 3x5 minutes with PBS and mounted using ProLong Glass Antifade Mountant (ThermoFisher). Images were taken on a Zeiss AxioObserver.A1 microscope.

### Comparative genomics

Two HAL alignments were used to assess conservation in differentially accessible regions: a 363-species avian alignment (https://cgl.gi.ucsc.edu/data/cactus/363-avian-2020.hal) and a 605-species mammalian+avian alignment (https://cgl.gi.ucsc.edu/data/cactus/605-vertebrate-2020.hal). For each alignment, the Bengalese finch assembly was replaced with the assembly used here (lonStrDom2) using the script *cactus-update-prepare* (https://github.com/ComparativeGenomicsToolkit/cactus/tree/master, v3.1.2). MAF files referenced to lonStrDom2 were exported from each HAL file using *cactus-hal2maf* (–outType single). The following approaches are described in full in Nextflow pipelines and configuration files available at https://github.com/colquittlab/song-system-grn. Neutral models were estimated using *phyloFit v.1.96* (https://github.com/CshlSiepelLab/ phast) on subsampled ancestral repeat elements. For each alignment, three models were estimated using three chromosome sets – macro (chr1-8), micro (chr9-29), and sex chromosomes (chrZ) – following the reasoning in [30] of different evolution rates across avian chromosomes. PhyloP was performed on each alignment using the three neutral models (mode = CONACC, method = LRT). For phastCons, we first estimated a scaling factor across 1 megabase alignment chunks then averaged the resulting conserved and non-conserved models within chromosome sets. PhastCons was then run using these models, an expected length of 20 bp, and an expected coverage of 0.25. Matrices for these tracks, as well as chromatin accessibility tracks, across differentially accessible regions were obtained using *deep-tools v3.5.4* [70] function *computeMatrix*.

### Large language models

Claude Sonnet 4.6 (Anthropic) was used in the following ways: to generalize analysis code (e.g. to loop through parameter combinations), to generate wrapper functions, to draft Nextflow pipelines, to troubleshoot errors, and to clean and document code.

### Data availability

The single-nucleus RNA-seq, Multiome-seq, and chicken overexpression datasets are available in the GEO repositories GSE316539, GSE316538, and GSE328195. A UCSC Genome Browser TrackHub containing ATAC tracks and conservation analysis is available at http://colquitt-track-hub.pbsci.ucsc.edu/lonStrDom2/hub.txt.

### Code availability

The publicly available data and code for analysis are available at https://github.com/colquittlab/song-system-grn (doi:10.5281/zenodo.19372037).

## Acknowledgments

This work was supported by the National Institutes of Health (NIH) Neurological Disorders and Stroke R01NS138781 to BMC and HHMI Investigator award to MSB. Sequencing was performed at the UCSF CAT (supported by UCSF PBBR, RRP IMIA, and NIH 1S10OD028511-01 grants) and the DNA Technologies and Expression Analysis Cores at the UC Davis Genome Center (RRID:SCR_017740, supported by NIH Shared Instrumentation Grant 1S10OD010786-01). The Zeiss 880 Confocal microscope is funded by NIH S10 Grant 1S10OD23528-01. The UCSC Life Sciences Microscopy Center (RRID:SCR_021135) is a core facility supported by the UCSC Division of Physical and Biological Sciences and the California Institute for Quantitative Biosciences (QB3).

## Author Contributions

A.M. performed snRNA-seq library preparation for the overexpression experiments and wrote the initial manuscript. C.L. performed histology assays and assisted in sequencing library preparations. E.K. performed *in ovo* electroporations of chick embryos. F.R. developed and optimized the protocol for *in ovo* electroporations of chick embryos. K.V. generated and validated the overexpression constructs. M.B. obtained funding for the project and edited the final draft. B.C. conceived of the project, acquired and analyzed snRNA-seq and Multiome-seq data, and wrote the initial draft.

## Supplementary Information

### Supplementary Discussion

#### Cell type labels compared to Colquitt, Merullo et al. 2021

Each nidopallial song region contained two projection neuron types (Glut-HVC-1, Glut-HVC-2, Glut-LMAN-1, and Glut-LMAN-2), while the arcopallial song region RA contained a single projection neuron type (Glut-RA). This is fewer than the number of HVC and RA glutamatergic neuron types described in our previous work [20]. We attribute this difference to two factors. First, this dataset consists of only a single species and data type (the previous dataset contained nuclei from zebra finches and cells from Bengalese finches), reducing batch effects that may have generated additional clusters. Second, the inclusion of surrounding tissue here allowed us to disambiguate whether clusters found in song regions are *bona fide* regional clusters or are the result of dissection error.

In HVC, several cell type label changes are worth specifically mentioning. Glut-HVC-2 here is identical to HVC-Glut-3 in [20] (**Extended Data Fig. 2c**), corresponding to HVC_X_ projection neurons. In addition, Glut-NC-1 and Glut-NC-2 are similar to HVC-Glut-2 in the previous analysis (**Extended Data Fig. 2c**). Although these types appear to be present in HVC (**Fig. 1f**), they are not specific to HVC and received the more general “NC” label. Similarly, HVC-Glut-5 in the 2021 analysis is labeled Glut-NC-3 here, given its much higher abundance in NC compared to HVC. Glut-HVC-1a lies intermediate to Glut-HVC-1 and Glut-NC-1 (**Fig. 1d**) and may represent cells transiting from the NC-1 state to the mature HVC-1 state.

#### LMAN projections

Like HVC, LMAN projects to both RA and Area X, yet current evidence indicates that these projections arise from a single projection population [71–73], leaving the projection identities of the two Glut-LMAN neuron types ambiguous. We propose that Glut-LMAN-2, which is transcriptionally similar to Glut-HVC-2 (**Fig. 2a**), is an uncharacterized population projecting to Area X (**Fig. 2b**).

### Supplementary Figures

**Extended Data Figure 1:**
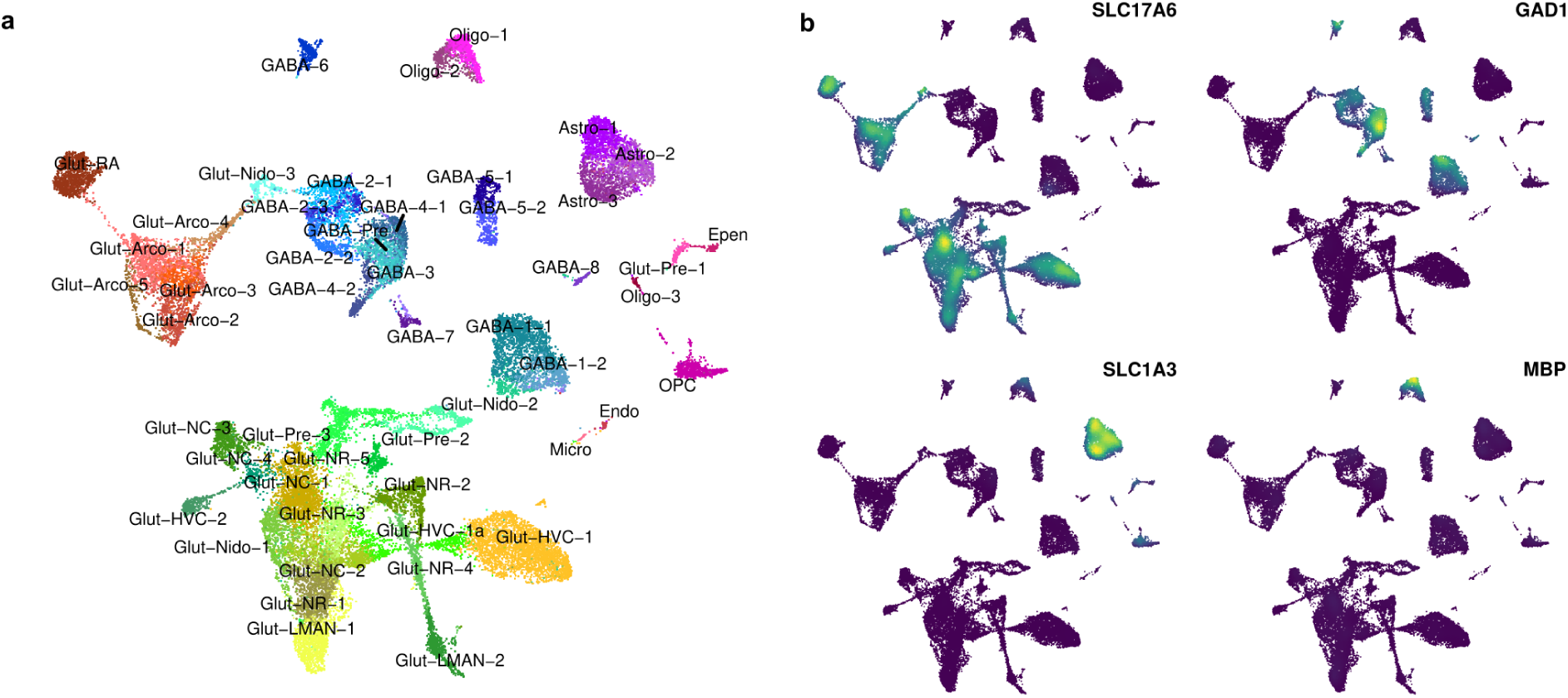
Overview of cell types across song and non-song regions. **(a)** UMAP plot of the snRNA-seq dataset showing all clusters. **(b)** UMAP plots showing the expression of *SLC17A6* (glutamatergic neurons), *GAD1* (GABAergic neurons), *SLC3A1* (astrocytes), and *MBP* (oligodendrocytes).

**Extended Data Figure 2:**
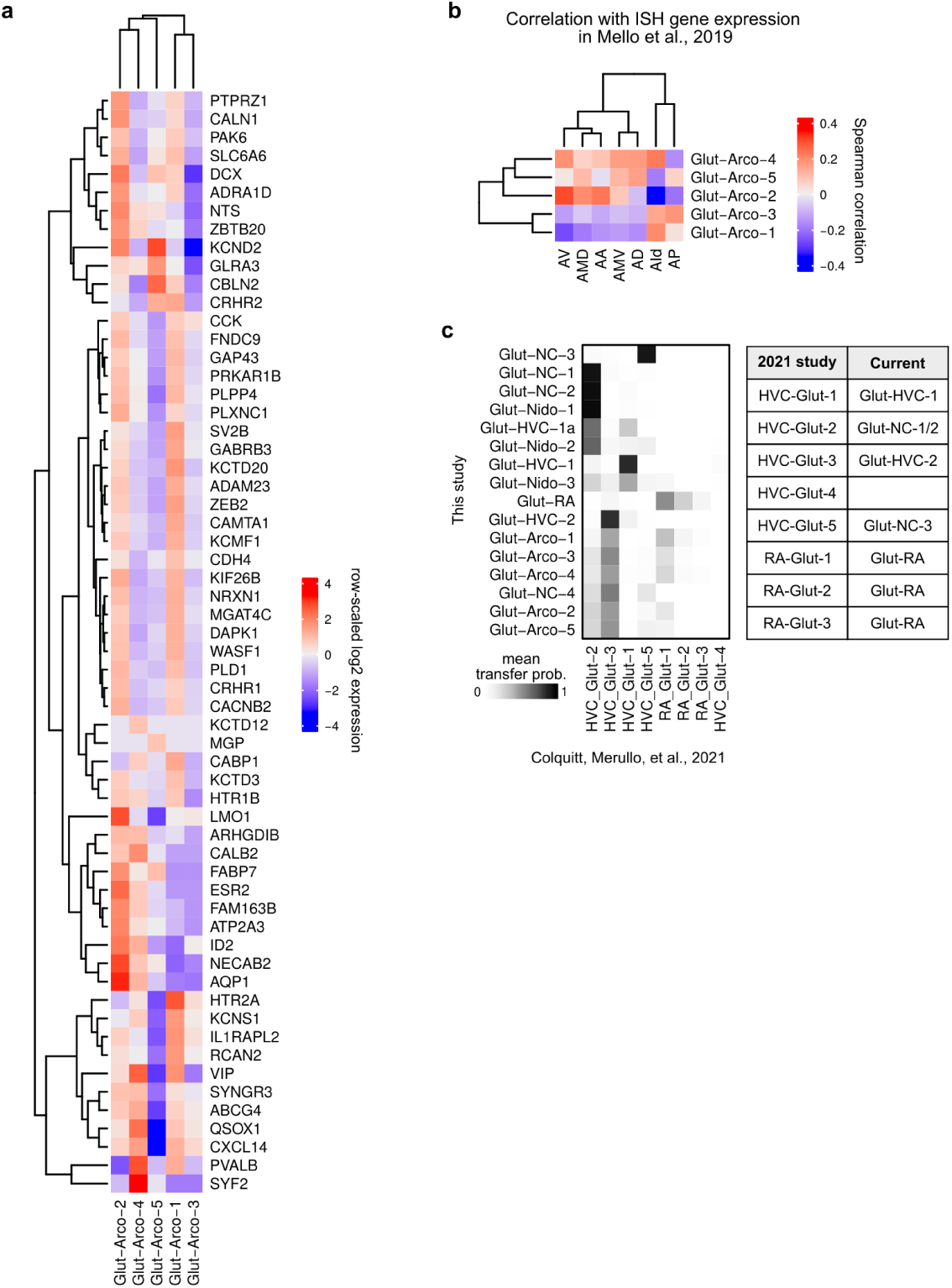
Transcriptional similarities of arcopallial glutamatergic neurons to arcopallial subregions. **(a)** Expression of the genes tested by *in situ* hybridization in [74] across the arcopallial glutamatergic populations described here. **(b)** Pairwise Spearman correlations of gene expression between arcopallial glutamatergic neuron types (snRNA-seq) and arcopallial regions (ISH). **(c)** Label transfer from [20] glutamatergic neuron labels to current glutamatergic neuron labels. (*Left*) Mean trans-fer probabilities from Seurat reciprocal PCA anchor-based integration. (*Right*) Table of label correspondences between the two studies.

**Extended Data Figure 3:**
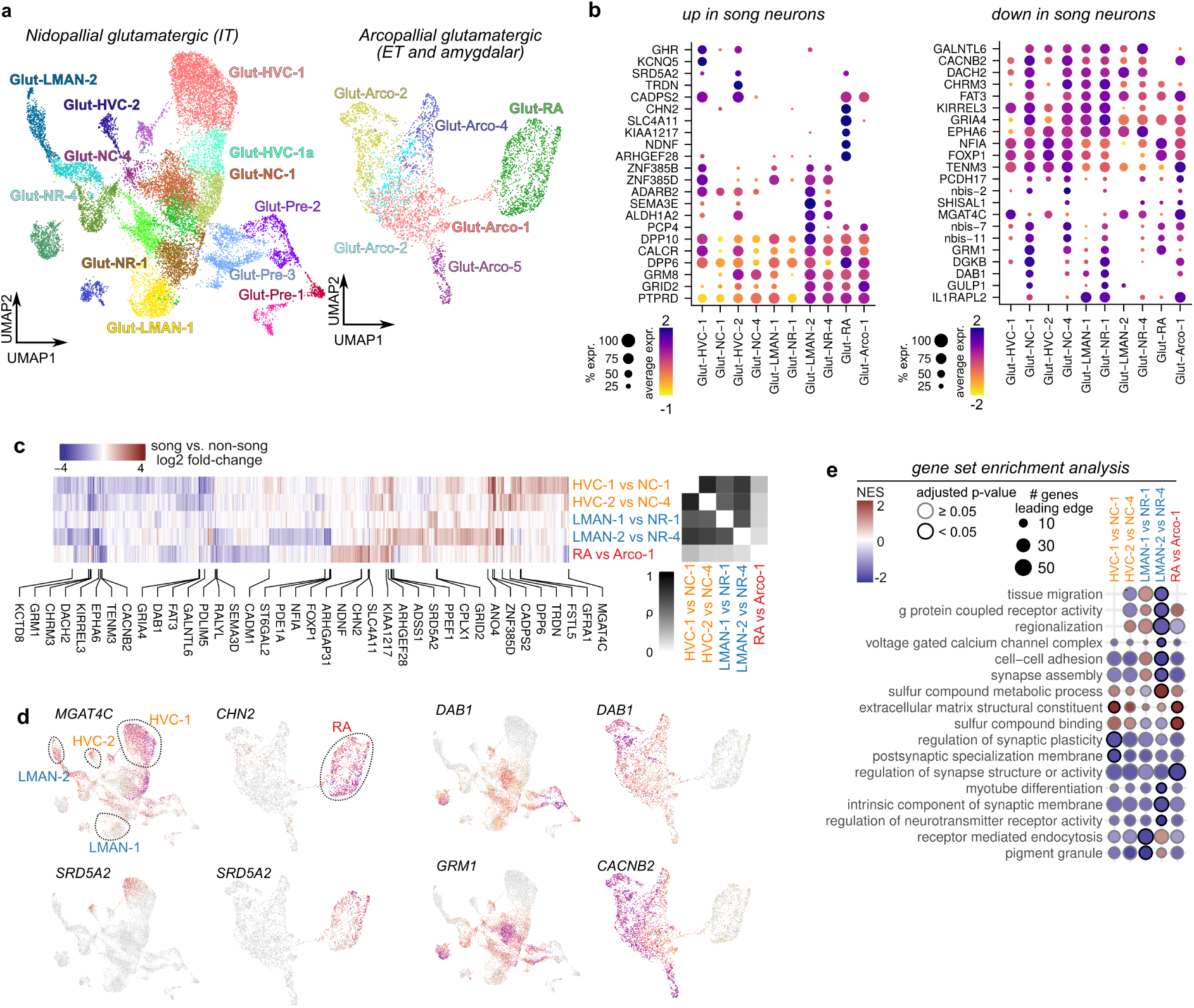
Molecular specialization of song glutamatergic neurons. **(a)** UMAP plot of glutamatergic neuron clusters from the (*left*) nidopallium, including HVC and LMAN, and (*right*) arcopallium, including RA **(b)** Expression of the top five differentially expressed genes (up or down) for each song vs non-song pair. **(c)** Heatmap of log2 fold changes in expression between each song glutamatergic neuron and its non-song counterpart. Grayscale heatmap shows the pairwise Spearman correlation of differential expression across each song neuron. **(d)** UMAP of nidopallial and arcopallial glutamatergic cells showing the expression of example genes that elevated or reduced expression in song neurons. **(e)** Gene set enrichment analysis of differentially expressed genes between each song and non-song glutamatergic neuron pair.

**Extended Data Figure 4:**
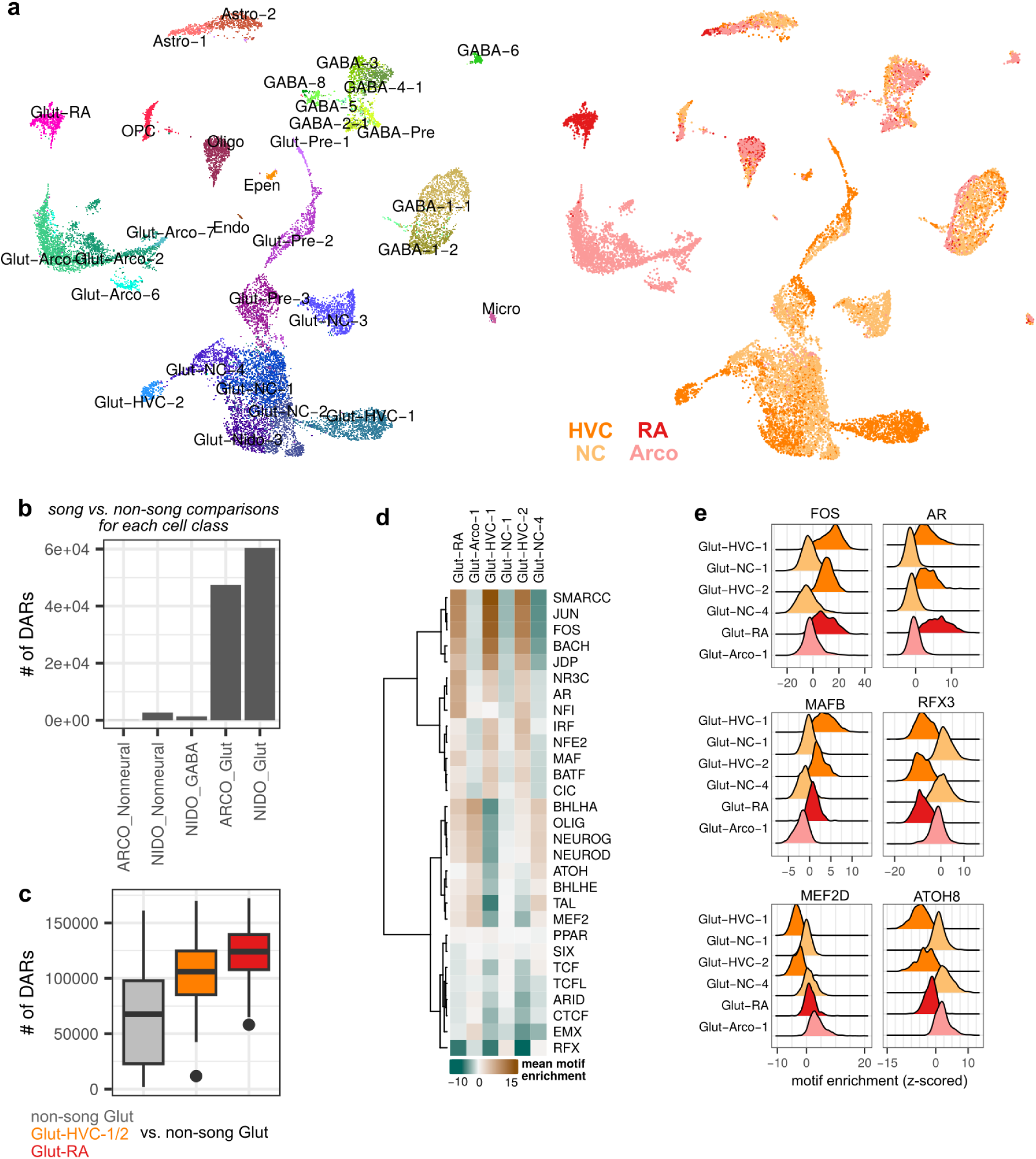
Enrichment of transcription factor motifs in song versus non-song glutamatergic neurons. **(a)** UMAP plot of the Multiome dataset, expanded to show (*left*) all cluster labels and (*right*) dissection regions **(b)** The number of differentially accessible regions (DARs) between song and non-song regions for each cell class. **(c)** Distributions of the numbers of DARs across pairs of glutamatergic neuron types. Comparisons are divided into three sets: (1) non-song Glut neurons versus non-song Glut neurons, (2) Glut-HVC1/2 versus nidopallial non-song Glut neurons, and (3) Glut-RA versus arcopallial non-song Glut neurons. **(d)** Mean enrichment of transcription factor binding site motifs across differentially accessible regions (DARs) in song and non-song glutamatergic types. Motifs are grouped by transcription factor family. **(e)** Selected motifs that are commonly enriched or depleted across the DARs of each song glutamatergic neuron type.

**Extended Data Figure 5:**
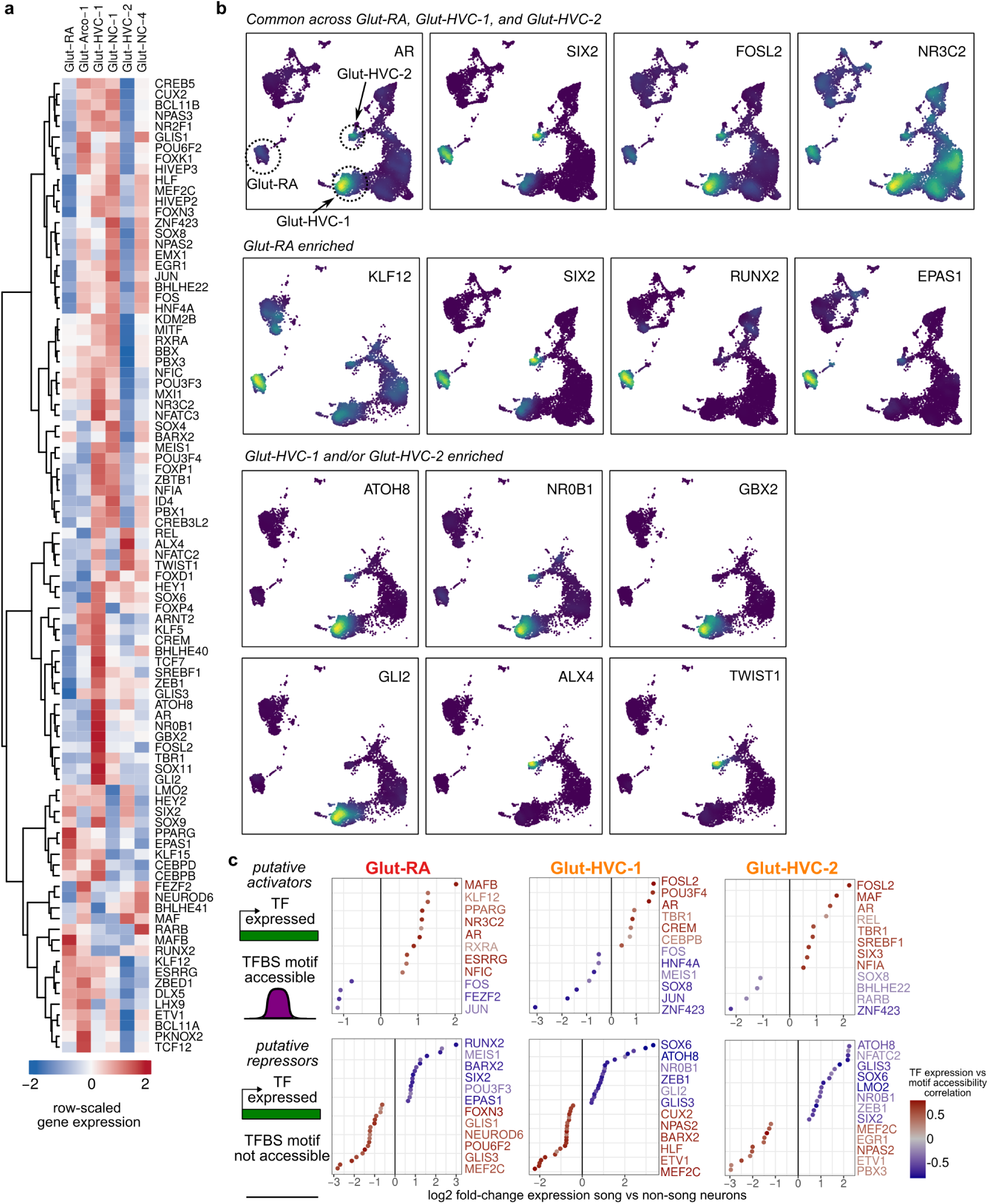
Expression of transcription factors differentially expressed across song glutamatergic neurons. **(a)** Heatmap of transcription factor gene expression in song glutamatergic neurons (Glut-RA, Glut-HVC-1, Glut-HVC-2) and paired non-song neurons (Glut-Arco-1, Glut-NC-1, Glut-NC-4). Expression of each gene is scaled across cell types. **(b)** UMAP plots of glutamatergic neurons showing the expression of selected transcription factors between song and non-song glutamatergic neurons. **(c)** The accessibility of TFBS motif-containing elements was correlated with the expression of associated transcription factors to identify putative transcriptional activators and repressors that are differentially expressed in song versus non-song glutamatergic neurons. At the right of each plot are transcription factors with the highest or lowest expression different between song and non-song neurons.

**Extended Data Figure 6:**
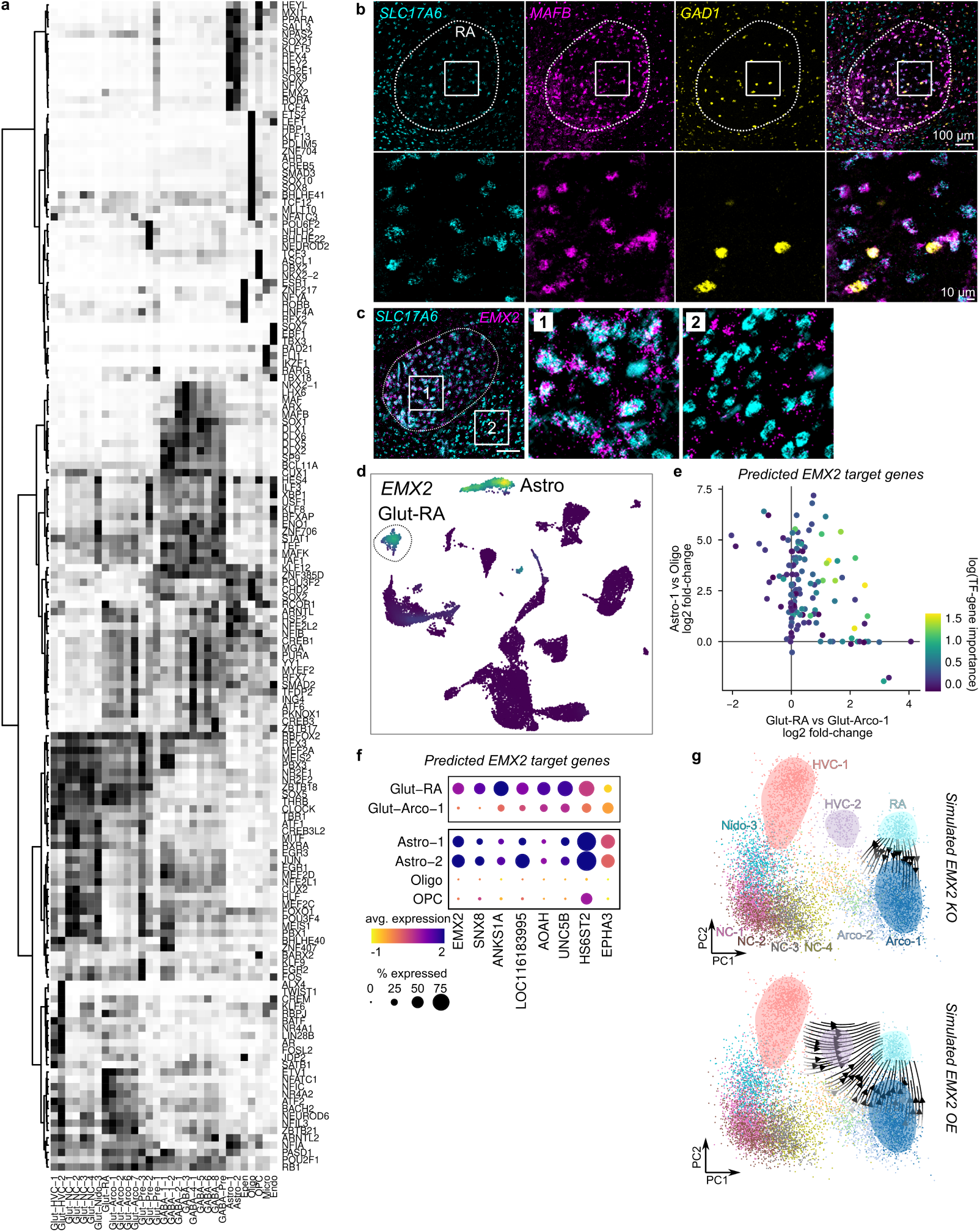
Gene regulatory network analysis of song glutamatergic neurons. **(a)** Expansion of heatmap in Figure 4a. **(b)** *In situ* hybridization indicating that MAFB is expressed in glutamatergic (*SLC17A6*-positive) and a subset of GABAergic (*GAD1*-positive) neurons in RA (outlined). However, in motor regions outside of RA that are not involved in song production, *MAFB* is only expressed in GABAergic neurons. **(c)** *EMX2* is expressed in glutamatergic neurons in RA but is not expressed in glutamatergic neurons outside of RA. Scale bar is 100 *µ*m. **(d)** UMAP plot of the *EMX2* expression. *EMX2* has elevated expression in Glut-RA neurons as well as in astrocytes (*EMX2*). **(e)** Differential expression (DE) of predicted *EMX2* target genes between Glut-RA and Glut-Arco-1 neurons compared to astrocytes (Astro-1) and oligodendrocytes (Oligo). Point color indicates TF to target gene importance from SCENIC+ tree-based regression. **(f)** Expression of selected genes showing elevated expression in Glut-RA and astrocytes. **(g)** Simulated (*top*) knockout and (*bottom*) overexpression of *EMX2* in glutamatergic neurons. Arrows indicate the direction of shifts in neuron position in PCA space following each manipulation.

**Extended Data Figure 7:**
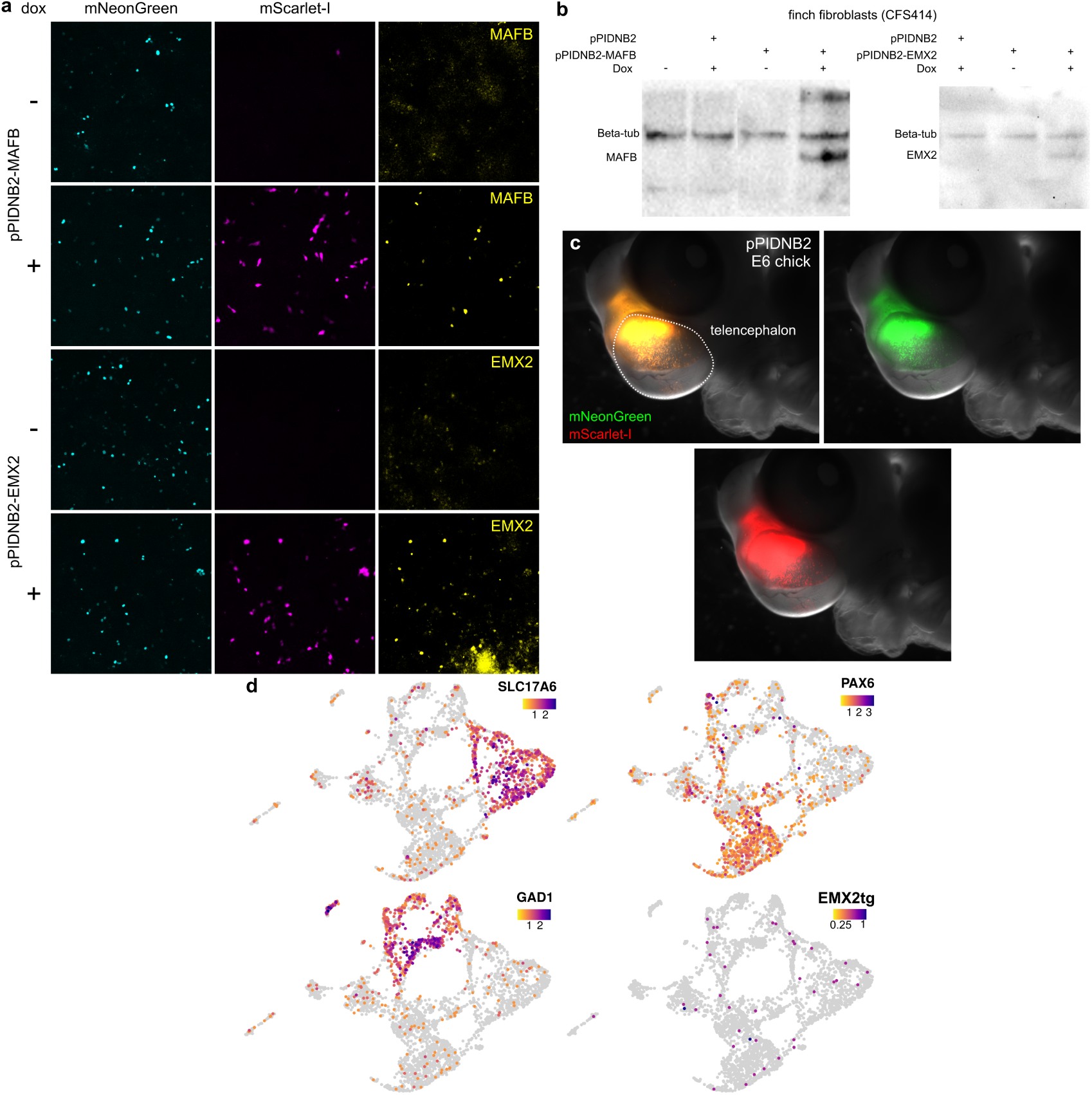
Validation of *MAFB* and *EMX2* overexpression construct. **(a)** Immunocytochemistry validation of dox-dependent expression of MAFB and EMX2 from pPIDNB2-MAFB and pPIDNB2-EMX2. **(b)** Western blot for MAFB and EMX2 overexpression in finch fibroblasts (CFS414) via the pPIDNB2-MAFB and pPIDNB2-EMX2 constructs, using *α*-MAFB, *α*-EMX2, and *α*-*β*-tubulin antibodies. **(c)** Expression of the pPIDNB2 construct in E6 chicken embryos following electroporation at E4 and doxycycline addition at E5. **(d)** UMAP plots showing the expression of *SLC17A6* (glutamatergic neurons), *GAD1* (GABAergic neurons), and *PAX6* (progenitor cells).

**Extended Data Figure 8:**
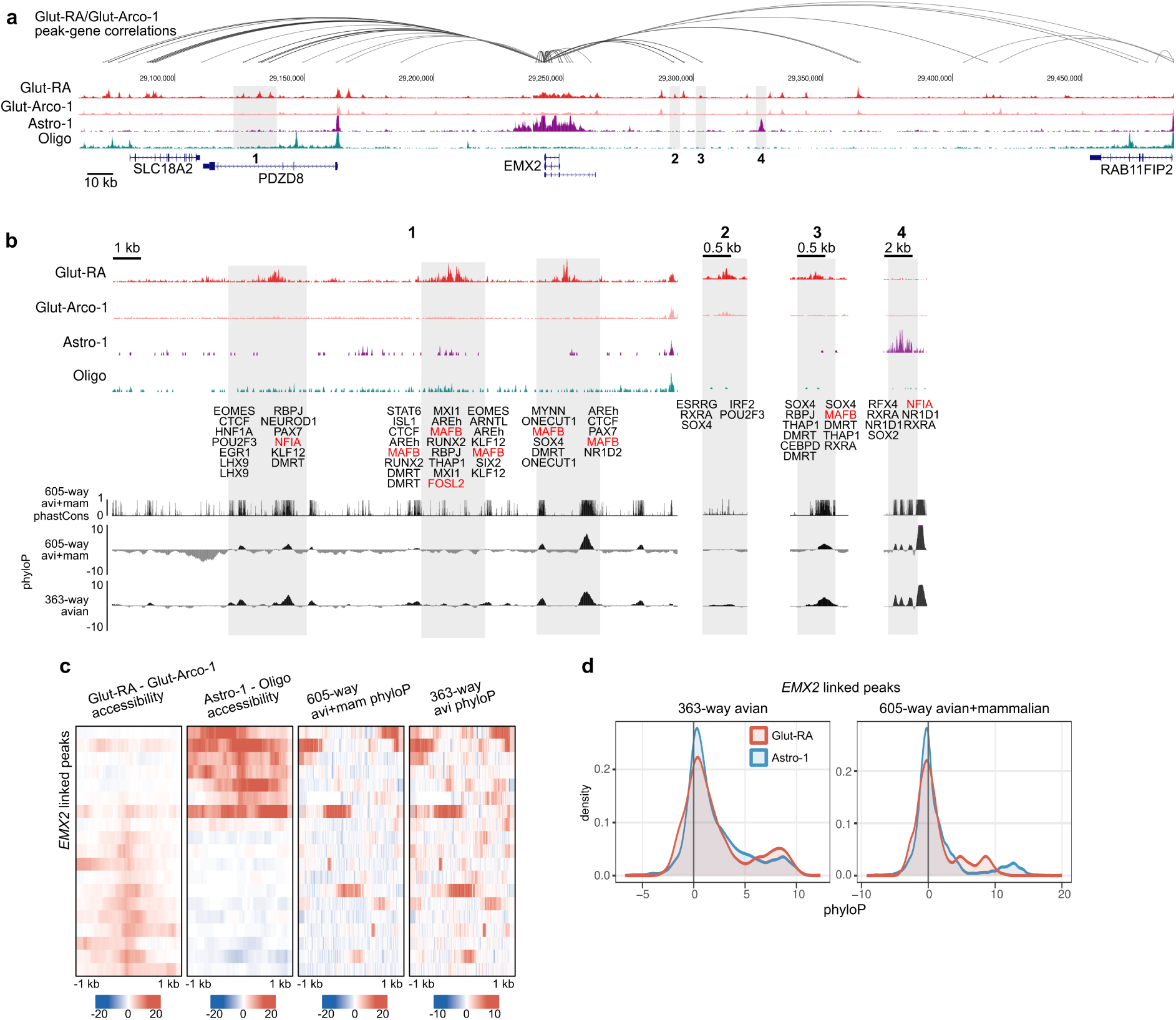
Song neuron-specific regulatory landscape of *EMX2*. **(a)** Chromatin accessibility flanking the *EMX2* locus. Arcs above the accessibility tracks link the MAFB transcription start site with DARs whose accessibilities covary with *EMX2* expression (Pearson r > 0.4). **(b)** Magnification of the numbered regions indicated in panel **b**. Transcription factor binding sites for each differentially accessible region are annotated. Sites for SCENIC+ predicted inputs are highlighted in red. AREh, androgen receptor element half-site; AREf, androgen receptor element full-site. At bottom are three tracks showing the conservation (phastCons) or conservation and acceleration (phyloP) across two alignments, one containing 605 species of birds and mammals (avi+mam) and a second with 363 species of birds only. **c)** Differential accessibility and phyloP scores across all DARs linked to *EMX2*. Distributions of phyloP scores across *EMX2*-linked DARs with high differential accessibility (normalized fragment density differences greater than 5 between Glut-RA and Glut-Arco-1 or Astro-1 and Oligo).

